# Broadly neutralizing antibodies targeting a conserved silent face of spike RBD resist extreme SARS-CoV-2 antigenic drift

**DOI:** 10.1101/2023.04.26.538488

**Authors:** Ge Song, Meng Yuan, Hejun Liu, Tazio Capozzola, Ryan N. Lin, Jonathan L. Torres, Wan-ting He, Rami Musharrafieh, Katharina Dueker, Panpan Zhou, Sean Callaghan, Nitesh Mishra, Peter Yong, Fabio Anzanello, Gabriel Avillion, Anh Lina Vo, Xuduo Li, Muzamil Makhdoomi, Ziqi Feng, Xueyong Zhu, Linghang Peng, David Nemazee, Yana Safonova, Bryan Briney, Andrew B Ward, Dennis R. Burton, Ian A. Wilson, Raiees Andrabi

**Affiliations:** Department of Immunology and Microbiology, The Scripps Research Institute, La Jolla, CA 92037, USA; IAVI Neutralizing Antibody Center, The Scripps Research Institute, La Jolla, CA 92037, USA; Consortium for HIV/AIDS Vaccine Development (CHAVD), The Scripps Research Institute, La Jolla, CA 92037, USA; Department of Integrative Structural and Computational Biology, The Scripps Research Institute, La Jolla, CA 92037, USA; Department of Computer Science, Johns Hopkins University, Baltimore, MD 21218, USA; Ragon Institute of Massachusetts General Hospital, Massachusetts Institute of Technology, and Harvard University, Cambridge, MA 02139, USA; Skaggs Institute for Chemical Biology, The Scripps Research Institute, La Jolla, CA 92037, USA; Lead Contact

## Abstract

Developing broad coronavirus vaccines requires identifying and understanding the molecular basis of broadly neutralizing antibody (bnAb) spike sites. In our previous work, we identified sarbecovirus spike RBD group 1 and 2 bnAbs. We have now shown that many of these bnAbs can still neutralize highly mutated SARS-CoV-2 variants, including the XBB.1.5. Structural studies revealed that group 1 bnAbs use recurrent germline encoded CDRH3 features to interact with a conserved RBD region that overlaps with class 4 bnAb site. Group 2 bnAbs recognize a less well-characterized "site V" on the RBD and destabilize spike trimer. The site V has remained largely unchanged in SARS-CoV- 2 variants and is highly conserved across diverse sarbecoviruses, making it a promising target for broad coronavirus vaccine development. Our findings suggest that targeted vaccine strategies may be needed to induce effective B cell responses to escape resistant subdominant spike RBD bnAb sites.

## Introduction

Broadly neutralizing antibody (bnAb) epitope-based vaccines are an important strategy for developing effective interventions against coronaviruses. Among the most potent and dominant neutralizing antibodies (nAbs) elicited in SARS-CoV-2 human infection or vaccination are those targeting the SARS-CoV-2 spike receptor binding domain (RBD) ^1–10^. Vaccines that induce these spike RBD nAbs have shown high effectiveness in reducing COVID-19 disease severity and hospitalization ^11–14^. However, the emergence of SARS-CoV-2 variants of concern (VOCs) has led to the majority of these RBD-targeting antibodies losing their neutralizing activity ^15–24^. The bulk of the mutations on the spikes of SARS-CoV-2 VOCs occur in the RBD region, resulting in substantially reduced potency or loss of neutralizing activity of most clinically approved RBD-targeting nAbs ^15, 21, 25^. This scenario highlights the urgent need to identify bnAbs that can target RBD epitopes that are more highly conserved and resistant to mutation. Such bnAbs are crucial for developing antibody-based interventions and variant-proof vaccines and may also be important against emerging coronaviruses with the potential to seed future pandemics in humans.

RBD nAbs have been classified into 4 major classes, class 1-4, and several subclasses ^3, 10, 26^. Class 1 and 2 RBD antibodies are the most potent and frequently elicited nAbs that target overlapping regions of the receptor binding site (RBS), where the host cell receptor ACE2 binds ^3, 6, 10, 26, 27^. These nAbs exhibit limited cross-reactivity with related coronaviruses and are easily escaped by SARS-CoV-2 variants ^26, 27^. Class 3 and 4 RBD nAbs are less potent and are less frequently elicited in humans but target relatively more conserved regions of the RBD and exhibit cross-reactivity with VOCs and diverse sarbecoviruses ^3, 26, 28–38^. Elicitation of nAb responses that target class 3 and 4 RBD sites or the nearby overlapping nAb epitopes are thus more desirable for broad sarbecovirus vaccine strategies.

In a previous study, we described two sets of RBD bnAbs, group 1 and group 2, that neutralize diverse ACE2-utilizing sarbecoviruses and exhibit binding to clade 2 and 3 non ACE2 sarbecovirus spike RBDs by targeting more conserved RBD epitopes ^34^. Group 1 RBD bnAbs are more potent in neutralization, while group 2 RBD bnAbs show relatively broader binding with different sarbecovirus clades, especially clade 2. Both group 1 and group 2 RBD bnAbs appear to be less frequently elicited in SARS-CoV-2 infection or vaccination and were isolated from two individuals with hybrid immunity (COVID-19 recovered and then vaccinated humans) ^34^.

In the current study, we investigated the molecular basis of sarbecovirus neutralization breadth by these group 1 and 2 RBD bnAbs and implications for broad vaccine strategies. We first tested the neutralization capacity of a select subset of the most potent and broadest group 1 and group 2 RBD bnAbs with recently emerged SARS-CoV-2 variants. We observed that some of these RBD bnAbs still retain neutralizing activities against highly evolved SARS-CoV-2 variants, including BA.4/5 and XBB.1.5. Group 2 RBD bnAbs were less affected by the more recent Omicron escape mutations. Furthermore, we determined crystal structures of multiple group 1 and group 2 RBD bnAbs to provide a molecular basis for the broad neutralization of sarbecoviruses and resistance to Omicron neutralization escape. The group 1 RBD bnAbs target a relatively conserved epitope proximal to the class 4 nAb target site or CR3022 cryptic site. The group 2 RBD bnAbs recognize a conserved and relatively ’silent’ face of the spike RBD, previously termed site V or lateral site ^10^. The group 2 RBD bnAb site V is cryptic on the native-like spike, and bnAbs targeting this site disrupt the spike as a possible mechanism of neutralization. Our data further suggest that both group 1 and 2 RBD bnAb memory B cells may be boostable with bivalent or heterologous SARS spike vaccines towards greater neutralization breadth. Overall, we provide a detailed molecular characterization of RBD bnAb epitopes that could serve as templates for the development of broad coronavirus vaccines, provided that appropriate immunogens can be engineered.

## Results

### Sarbecovirus spike RBD bnAbs that resist SARS-CoV-2 antigenic escape

As SARS-CoV-2 antibody escape variants continue to emerge, it has become increasingly important to identify bnAbs targeting conserved epitopes that can tolerate the large number of antigenic escape mutations found especially on the Omicron variants. We previously isolated a broad panel of sarbecovirus bnAbs from COVID-19-recovered donors who were subsequently vaccinated ^34^. These bnAbs target two distinct regions on the RBD, categorized as group 1 and group 2, based on competition binning studies using SARS-CoV-2 nAbs of known specificities. Group 1 bnAbs compete with RBD class 4 site nAbs and group 2 target a less well-characterized conserved RBD region. Here, to investigate the ability of these bnAbs to resist SARS-CoV-2 escape mutations, we tested the neutralization ability of the group 1 (n = 14) and group 2 (n = 5) RBD bnAbs against a broad panel of SARS-CoV-2 variants including Omicron lineage variants (Figure 1 and Table S1). Group 1 RBD bnAbs were found to be relatively more potent and neutralized the early SARS-CoV-2 variants (Alpha, Beta, Gamma, and Delta) equally efficiently; however, their neutralizing activities dropped substantially against the Omicron variants (Geometric mean IC_50_ change: range = 14 – 105-fold IC_50_ drop) (Figure 1B and Table S1). The most pronounced neutralization loss was observed against BA.2 (geometric mean IC_50_ drop = 105-fold) and BA.2.75 (geometric mean IC_50_ drop = 104-fold) Omicron variants, respectively. Notably, four of the group 1 RBD bnAbs (CC25.3, CC25.36, CC25.54 and CC84.24) retained neutralizing activity (albeit less potently with Omicron variants) with most or all of the SARS-CoV-2 variants tested (Table S1).

**Figure 1.**
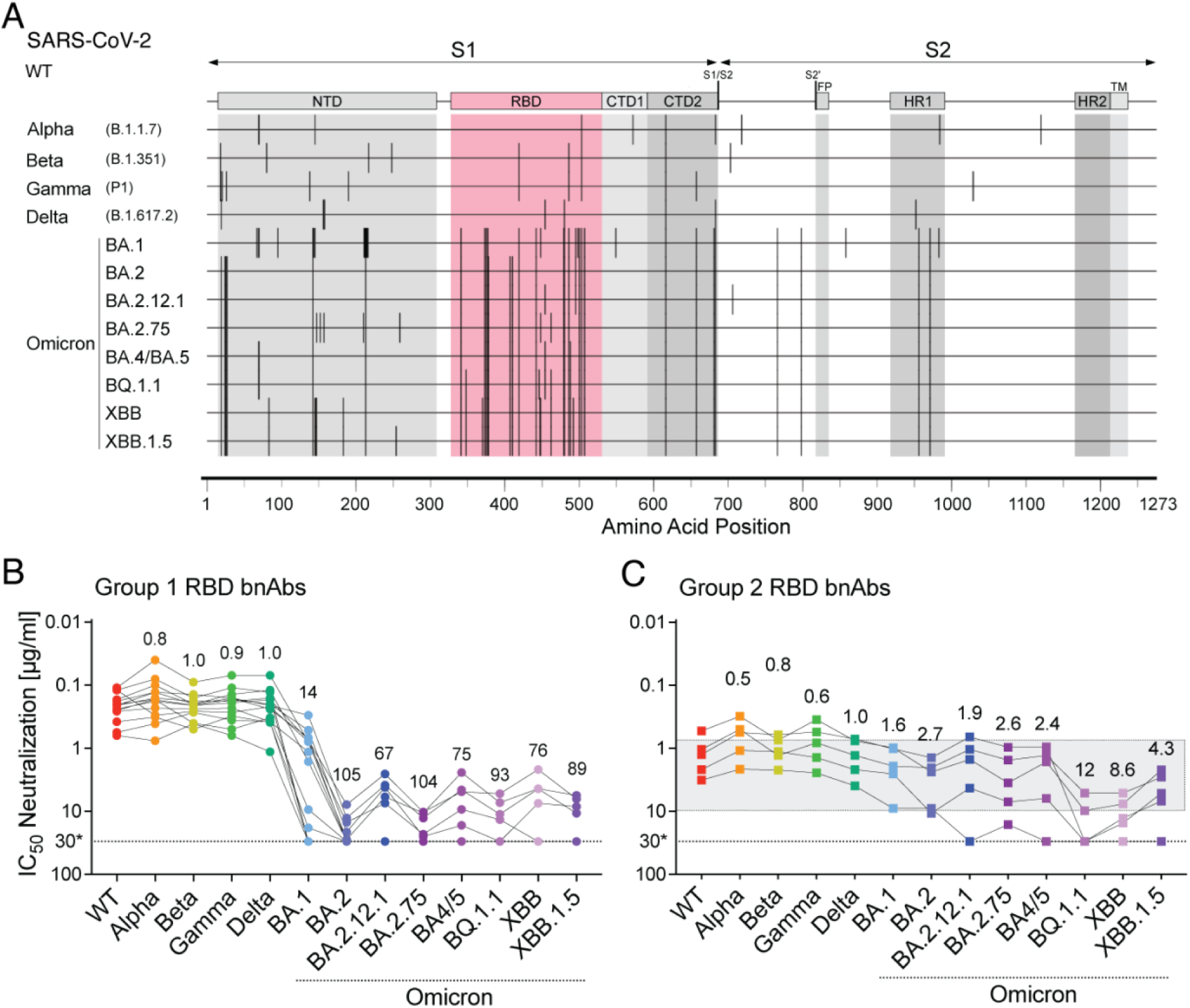
Neutralization of SARS-CoV-2 variants by RBD human broadly neutralizing antibodies. **A.** Schematic showing amino acid sequence alignment of SARS-CoV-2 spike region across 12 VOCs including Alpha (B.1.1.7), Beta (B.1.351), Gamma (P.1), Delta (B.1.617.2), Omicron subvariants BA.1, BA.2, BA.2.12.1, BA.2.75, BA.4/BA.5, BQ.1.1, XBB, and XBB.1.5. Vertical lines indicated mismatches compared to SARS-CoV-2 WT (Wuhan). Amino acid positions are illustrated at the bottom, S1 and S2 subunits are shown by arrowed lines at the top. Different domains are indicated (NTD, N-terminal domain; RBD, receptor-binding domain; CTD1, C-terminal domain 1; CTD2, C-terminal domain 2; S1/S2, S1/S2 furin cleavage site; S2′, S2′ TMPRSS2 or cathepsin B/L cleavage site; FP, fusion peptide; HR1, heptad repeat 1; HR2, heptad repeat 2; TM, transmembrane anchor), with the RBD region highlighted in red, containing the majority of the mutations. **B-C.** Neutralization IC_50_ of RBD bnAbs from group 1 (**B**) and group 2 (**C**) for the panel of SARS-CoV-2 WT and variants. Each variant is indicated by a different color. Round data points represent group 1 RBD bnAbs, square data points represent group 2 RBD bnAbs. The dotted line shows the limit of antibody dilution (30µg/ml). Values above each column indicate fold change of IC_50_s against each variant in comparison with WT. The grey shaded area highlights the bnAbs in group 2 showing relatively consistent neutralization IC_50_ across SARS-CoV-2 WT and variants.

Group 2 RBD bnAbs were shown to have an intrinsically lower neutralization IC_50_ potency, but were comparatively less sensitive to Omicron variant mutations (Geometric mean IC_50_ change: range = 2 – 17-fold IC_50_ drop) (Figure 1C and Table S1). Similar to group 1 RBD bnAbs, neutralization by the group 2 RBD bnAbs was minimally affected against the non Omicron SARS-CoV-2 variants. One out of five group 2 RBD bnAbs substantially lost neutralization with the Omicron variants (Table S1), but 4 of the 5 group 2 RBD bnAbs (CC25.4, CC25.17, CC25.43 and CC25.56) retained neutralization activities, albeit with IC_50_’s in the μg/ml range, against most or all of the variants tested. The IC_50_ neutralization changes for these 4 group 2 RBD bnAbs against the SARS-CoV-2 Omicron variants were modest suggesting targeting of spike epitopes that are relatively more resistant to antibody immune escape. The BQ.1.1 variant displayed the most neutralization resistance to group 2 RBD bnAbs compared to the WT SARS-CoV-2 (12-fold drop in geometric mean IC_50_). Nevertheless, the neutralization activities of 4 group 2 RBD bnAbs remained largely unchanged against the XBB.1.5 variant (Table S1), which is the most dominant SARS-CoV-2 variant (85% infections) circulating in the United States as of April 2023. As a comparison, we also tested 5 RBD nAbs that have been shown to target conserved RBD epitopes ^30, 32, 36, 39, 40^. Except for class 3 RBD site bnAb, S309, which retained neutralization against the Omicron variants, all other nAbs lost neutralization with these highly resistant SARS-CoV-2 variants (Table S1).

Altogether, we noted that both group 1 and 2 RBD bnAbs, and particularly the group 2 RBD bnAbs can effectively resist the extreme Omicron lineage antigenic drift and represent examples of human bnAbs that still retain substantial neutralizing activity with these highly evolved SARS-CoV-2 variants. These features support the potential utilization of group 1 and 2 RBD bnAbs in antibody-based interventions and as templates for variant-proof SARS-CoV-2 vaccines.

### Somatic hypermutation in RBD bnAbs is critical for neutralization of Omicron lineage variants

To investigate the role of antibody somatic hypermutation (SHM) in virus neutralization by group 1 and 2 RBD bnAbs, we generated inferred germline (iGL) versions by reverting their heavy and light chains to the corresponding germlines. The iGL heavy and light chain V, D and J regions were reverted back to their germline genes, while the non-templated N-additions at V/(D)/J junctions remained the same as in the mature bnAb versions, as described previously ^41^ (Figure 2A). It is not possible to revert the non-templated CDR3 junctional regions in the iGLs and these regions may potentially contribute to neutralization. Therefore, the iGLs primarily allow us to assess the contribution of the SHMs in the templated V-D-J regions for neutralization. We evaluated the mature group 1 and 2 RBD bnAbs and their iGL versions against SARS-CoV-2 variants. We noted that while many of the group 1 RBD bnAb iGLs retain neutralization with less mutated SARS-CoV-2 variants (Alpha, Beta, Gamma and Delta), they fail to neutralize the more evolved Omicron variants (Figure 2B and Table S1). In comparison the group 2 RBD bnAb iGLs fail to neutralize any of the SARS-CoV-2 variants (Figure 2C and Table S1). We also tested these group 1 and group 2 RBD bnAbs and their iGLs with ACE2-utilizing clade 1b (Pang17) and clade 1a (SARS-CoV-1 and WIV1) sarbecoviruses ^42, 43^. We noted that, while most mature bnAbs neutralized these sarbecoviruses, as reported previously ^34^, some iGLs of group 1 RBD bnAbs retained neutralization, especially with WIV1 and Pang 17 (Table S1). These results suggest that human antibodies in their germline configurations are able to recognize these sarbecovirus spikes, as also noted by other studies ^44^.

**Figure 2.**
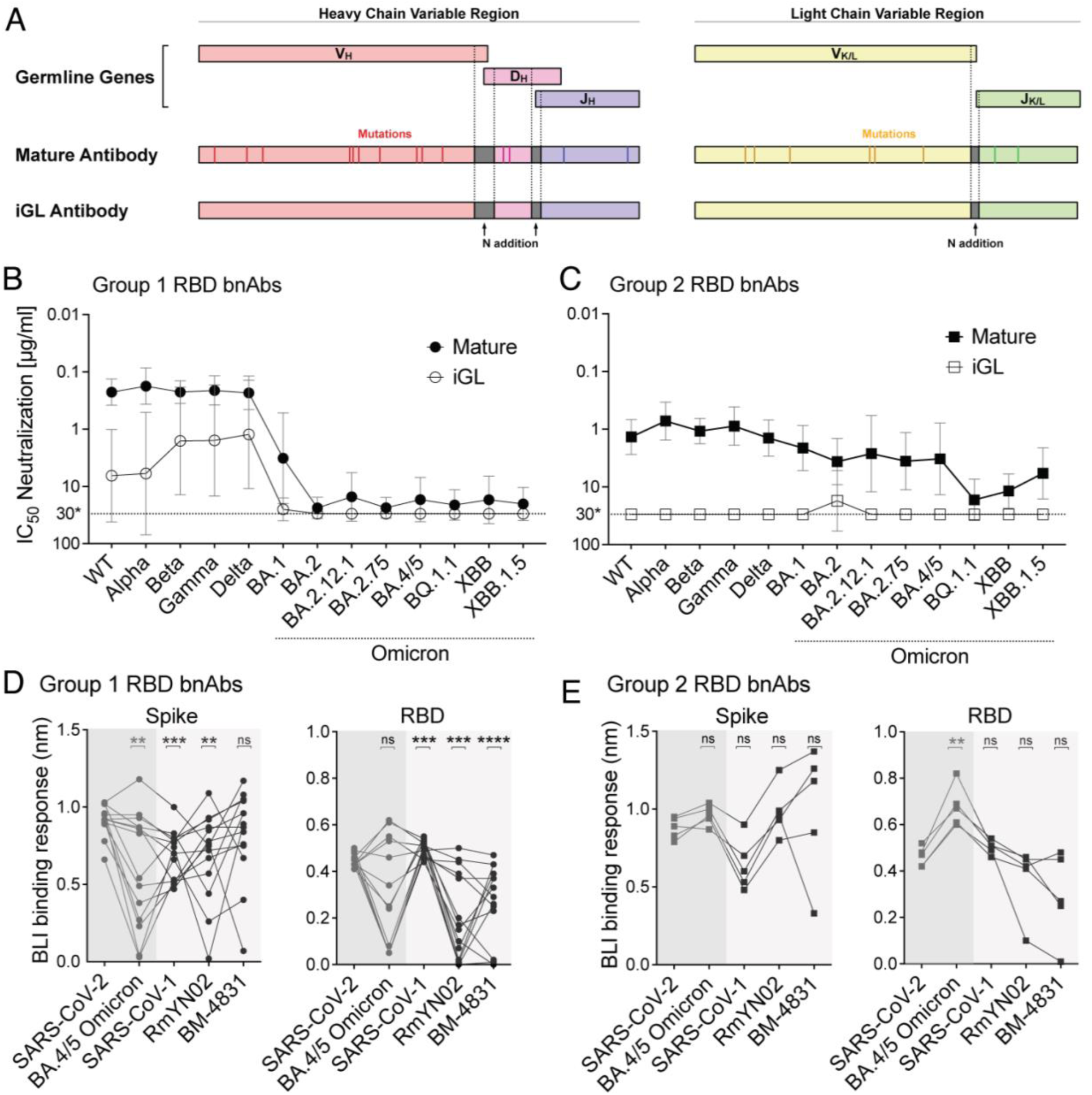
Neutralization and binding of mature RBD bnAbs and their inferred germline (iGL) versions against SARS-CoV-2 VOCs and other sarbecoviruses. **A.** Schematic showing design of inferred germline versions of bnAbs based on mature antibody heavy chain (left) and light chain (right) variable region sequences. The V/D/J genes of heavy chains (V_H_, D_H_, J_H_) and V/J genes of kappa or lambda light chains (V_K/L_, JK/L) were reverted to their corresponding germlines (IMGT/V-QUEST tool). The mutations represented by vertical lines were eliminated in the inferred germline antibody sequences. The non-templated N additions between V(/D)/J, indicated as dark grey, remained the same as in the mature antibody. **B-C.** Average neutralization IC_50_ values of all group 1 (B) and group 2 (C) RBD bnAbs comparing the mature antibody (bold) to their inferred germ line versions (open) tested with pseudotyped versions of SARS-CoV-2 WT (Wuhan) and 12 VOCs including Alpha (B.1.1.7), Beta (B.1.351), Gamma (P.1), Delta (B.1.617.2), Omicron subvariants BA.1, BA.2, BA.2.12.1, BA.2.75, BA.4/5, BQ.1.1, XBB, and XBB.1.5. Round data points represent group 1 RBD bnAbs, square data points represented group 2 RBD bnAbs. Each data point represents the geometric mean ± geometric SD of neutralization IC_50_ for specific variants by all bnAbs within the corresponding group (n = 14 for group 1, n = 5 for group 2). **D-E.** BLI binding response (nm) of group 1 (D) and group 2 (E) RBD bnAbs to the trimeric stabilized spike proteins and monomeric RBD proteins of SARS-CoV-2 (Wuhan and BA.4/5), clade 1b SARS-CoV-1, clade 2 RmYN02, and clade 3 BM-4831 sarbecoviruses. Statistical comparisons between groups were performed using a two-tailed Mann Whitney U-test (ns: p >0.05, *: p < 0.05, **: p < 0.005, ***: p < 0.001, ****: p < 0.0001).

Overall, the findings suggest that germline residues in group 1 RBD bnAbs may contribute to neutralization of SARS-CoV-2 and its minimally mutated variants. However, SHMs for both group 1 and 2 RBD bnAbs are required to neutralize more evolved SARS-CoV-2 Omicron lineage variants.

### Group 1 and 2 RBD bnAb memory B cells and potential recall boosts

To investigate whether group 1 and 2 RBD bnAbs bind to SARS-CoV-2 Omicron BA.4/5 spike and other clades of sarbecovirus spikes for potential boost considerations, we tested their binding to various spikes and their corresponding RBDs by BioLayer Interferometry (BLI). The BA.4/5 Omicron spike is a component of the current SARS-CoV- 2 bivalent booster vaccines ^45, 46^. Therefore, we first assessed whether the group 1 and 2 RBD bnAbs can bind to the BA.4/5 Omicron spike and whether their memory B cells are likely to be boosted by the current bivalent vaccines. As expected from the neutralization results, both group 1 and 2 RBD bnAbs showed strong binding to the SARS-CoV-2 spike protein (Figure 2D-E and Table S2). Consistent with the neutralization of BA.4/5 Omicron variant above, the binding of group 1 RBD bnAbs was significantly reduced against the BA.4/5 Omicron spike (Figure 2D). Nevertheless, many group 1 RBD bnAbs, especially the ones that neutralize BA.4/5 variant still bound to its spike protein with high affinity (Figure 2D and Table S2). In comparison, the group 2 RBD bnAbs bound to BA.4/5 Omicron spike equally efficiently (Figure 2E and Table S2). The findings suggest that majority of both group 1 and 2 RBD bnAbs are likely to be boosted with the BA.4/5 bivalent vaccine.

We further tested the binding by group 1 and 2 RBD bnAbs to spike proteins derived from heterologous clade 1a (SARS-CoV-1), clade 2 (RmYN02) and clade 3 (BM-4831) sarbecoviruses ^34, 42, 47^. The group 1 RBD bnAb showed substantially reduced binding to SARS-CoV-1 and RmYN02 compared to the SARS-CoV-2 spike but the binding with sarbecovirus clade 3 BM-4831 spike was comparable (Figure 2D and Table S2). Most of the group 2 RBD bnAbs showed strong binding to the clade 2 (RmYN02) and clade 3 (BM-4831) sarbecovirus spike-derived proteins (Figure 2E and Table S2). The results suggest that heterologous clades 2 and 3 spike-derived protein immunogens could be utilized to boost group 1 and 2 RBD bnAb responses, and specifically the clade 3 BM- 4831 spike immunogen may recall group 1 RBD bnAb memory B cells more efficiently. We also tested the group 1 and 2 RBD bnAbs with RBD of SARS-CoV-2, BA.4/5 Omicron and the heterologous clade 1a, 2 and 3 sarbecoviruses (SARS-CoV-1, RmYN02 and BM-4831). The binding responses were overall lower but largely consistent with corresponding spike binding with some exceptions (Figure 2D-E and Table S2).

Overall, we observed that the binding of group 1 RBD bnAbs with BA.4/5 Omicron and heterologous sarbecovirus spikes was substantially reduced as compared to the parental SARS-CoV-2 but binding by group 2 RBD bnAbs were comparable. Nevertheless, both groups of bnAbs are likely boostable by these spikes.

To gain more insight into the detailed binding modes of group 1 and 2 bnAbs, we selected four antibodies from group 1 and three antibodies from group 2 to perform detailed structural studies as described below.

### Structures of group 1 bnAbs complexed to SARS-CoV-2 RBD show a recurrent YYDRxG feature in CDRH3 and diverse light chain interactions

Six of the 14 antibodies from group 1 RBD bnAbs shared the same YYDRxG motif encoded by IGHD3-22 that we and others previously identified ^34, 48, 49^. These YYDRxG antibodies exhibited broad neutralization breadth against SARS-CoV-2 variants, including Omicron subvariants (Table S1). Two additional antibodies featuring a YYDSSG motif, a germline precursor of the YYDRxG motif ^48^, showed cross-reactive binding to, but no neutralization of, Omicron variants.

To understand the basis for the superior breadth of YYDRxG antibodies, we determined crystal structures of three antibodies, CC25.54, CC84.24, and CC84.2, in complex with SARS-CoV-2 RBD with resolutions ranging from 2.9 to 3.1 Å (Figure 3A-C and Table S3). The structures revealed that the antibodies bind the CR3022 cryptic site using similar approach angles that allow them to compete with ACE2 binding even through there is no or minor epitope overlap (Figure 3D-E and Figure S1). The approach angle is also similar to that identified previously for YYDRxG antibodies ^10, 48, 49^. Analysis of the buried surface area (BSA) on SARS-CoV-2 RBD by these antibodies using the PISA program found similar overall BSA, although the percentage of light chain BSA varies among different antibodies (Figure 3F and Figure S1). Further inspection of the antibody-antigen interactions showed variable contacts of the antibody light chains with SARS-CoV-2 RBD, while the heavy chain CDR3 maintained essentially the same contacts. The light chain of CC84.2 contributes a larger BSA compared to CC25.54 and CC84.24, involving CDRs L1 and L2 of the antibody (Figure 3F and Figure S2). Different germline genes encoding the light chains of these YYDRxG antibodies are responsible for the different interactions. CC84.2 is encoded by IGKV3-20, while CC25.54 and CC84.24 are encoded by IGLV3- 21 and IGKV1-5, respectively.

**Figure 3.**
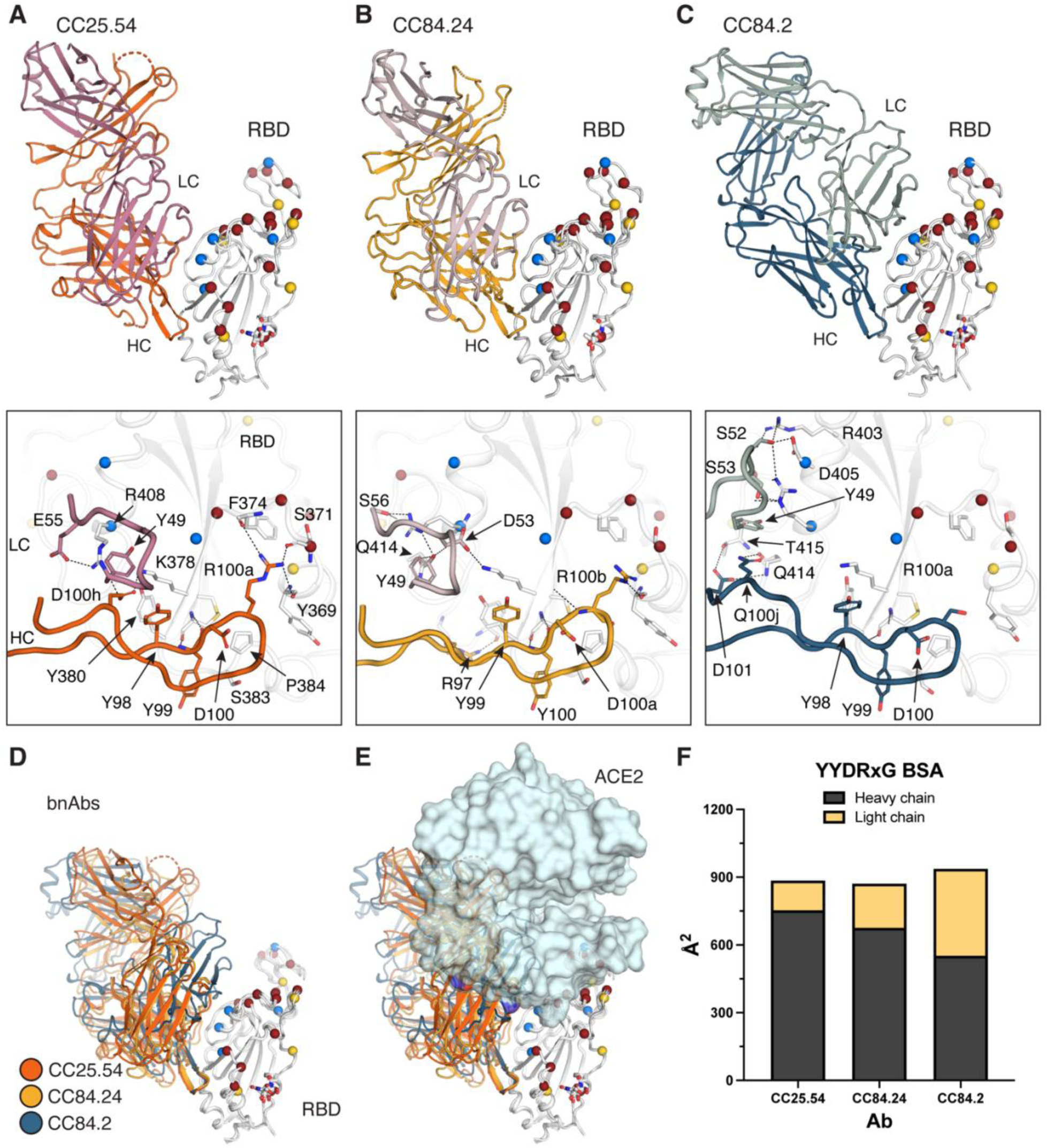
Crystal structures of representative YYDRxG antibodies from group 1 bnAbs. X- ray structures of two YYDRxG antibodies, CC25.54 (**A**) and CC84.24 (**B**), and a YYDSSG (YYDRxG precursor) antibody CC84.2 (**C**) are shown in ribbon representation (sticks represents observed N343 glycans in SARS-CoV-2 RBD crystal structure). The same perspective view is used for easy comparison. The CC25.54 heavy chain is colored in orange, light chain in rose pink; CC84.24 heavy chain in yellow, light chain in violet; CC84.2 heavy chain in navy blue, and light chain in teal. Residues from CDRH2 and H3 that interact with the RBD are shown in sticks. Dashed lines represent polar interactions. RBD residues that are mutated in Omicron are shown as spheres. Red represents residues mutated in BA.1, and additional mutations in blue in BA.4/5 and yellow in XBB.1.5. (**D**) Overlay comparison of YYDRxG antibodies determined in this study. Crystal structures are superimposed on SARS-CoV-2 RBD to compare the approach angles of these antibodies. (**E**) YYDRxG antibodies clash with ACE2 binding, although their epitope footprints do not overlap. Composite structures of YYDRxG antibodies from this study and ACE2 in complex with SARS-CoV-2 RBD (PDB ID: 6M0J) were used for comparison. (**F**) Comparison of the buried surface area (BSA) on SARS-CoV-2 RBD from the heavy and light chains of YYDRxG antibodies.

### A YYDML motif enables group 1 RBD bnAb CC25.36 to bind SARS-CoV-2 RBD in a similar approach angle as YYDRxG antibodies

Another group 1 antibody, CC25.36, showed comparable breadth as the YYDRxG antibodies. The crystal structure of CC25.36 revealed that the antibody binds SARS-CoV-2 RBD at the same CR3022 cryptic site on SARS-CoV-2 RBD, which appears less sensitive to mutations in Omicron strains (Figure 4A, Tables S1 and S2). Structural overlay illustrated that CC25.36 uses a similar approach angle to YYDRxG antibodies (Figure 4D). Further inspection showed that a YYDML motif in CC25.36 binds to approximately the same site as the YYDRxG motif in the other antibodies but with different interactions (Figure 4B and Figure S2). The _99_YY_100_ dipeptide in the YYDML motif binds with similar interactions as in the YYDRxG motif, which probably determines the site specificity of antibody binding (Figure 4B); the other RBD interactions with CDRH3 differ from the YYDRxG antibodies. D100a forms internal hydrogen bonds with the backbone of _100b_ML_100c_ as well as SARS-CoV-2 RBD K378. _100b_ML_100c_ interacts with a hydrophobic patch formed by RBD Y369, F374, and F378. PISA analysis confirmed that the light chain of CC25.36 contributes a large BSA similar to CC84.2 (Figure 4C and Figure S2). Structural superimposition on SARS-CoV-2 RBD showed that the CC25.36 light chain is positioned in the same way as CC84.2, although the CC25.36 light chain is encoded by the IGLV1-40 gene. A homology search for the YYDML motif showed that YYDIL or YYDLL motifs, also encoded by IGHD3-9, are present in other SARS-CoV-2 antibodies, e.g., COV2-2258 ^8^, C531 ^50^, and C2179 ^51^. However, whether they bind to the same epitope as CC25.36 warrants further investigation by structural studies.

**Figure 4.**
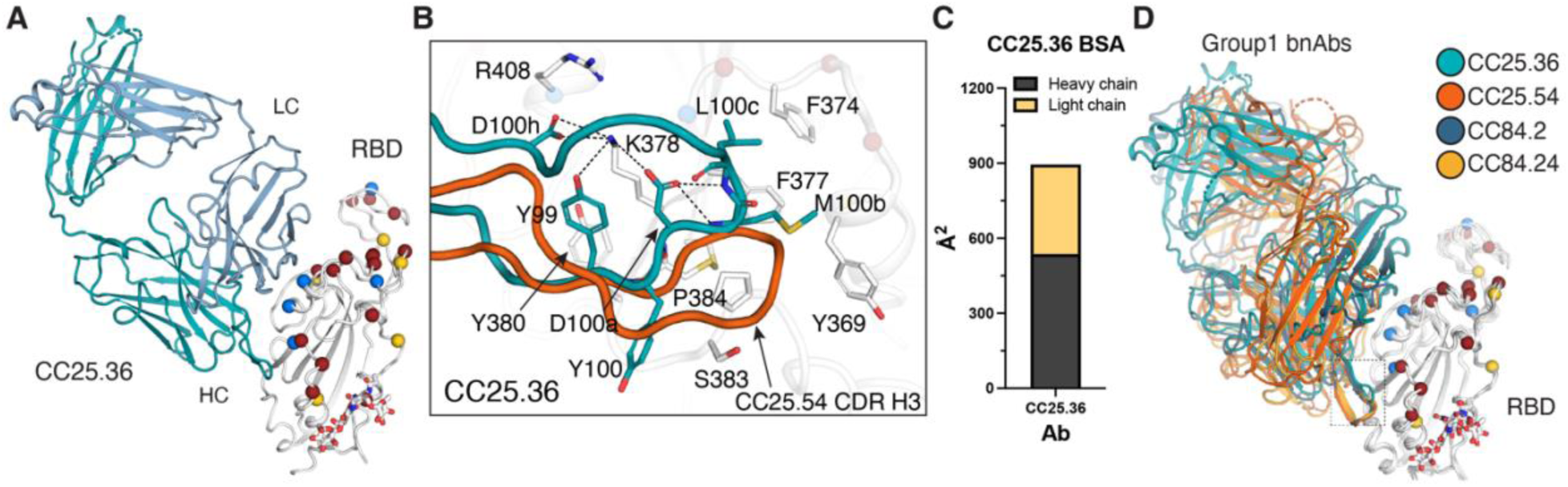
Crystal structure of group 1 antibody CC25.36 in complex with SARS-CoV-2 RBD. Heavy chain is colored in turquoise blue, and light chain in baby blue. RBD residues that are mutated in Omicron are shown as spheres. Red represents those mutated in BA.1; blue, additional mutations in BA.4/5; yellow, additional mutations in XBB.1.5. **A.** Overall structure of CC25.36 is shown in ribbon representation. Sticks represent glycans at N343 in SARS-CoV-2 RBD. **B.** Interactions between CDRH3 and RBD. Residues involved in the antibody-antigen interface are shown in sticks. The same perspective view as Figure 3A was used. CC25.54 CDR H3 was superimposed for easy comparison with Fig. 3 in orange. **C.** BSA on SARS-CoV-2 RBD from the CC25.36 heavy and light chains. **D.** CC25.36 CDRH3 uses a distinct binding mode. Structures of CC25.36 and YYDRxG antibodies determined in this study are superimposed on the RBD. CC25.36 CDR H3 (boxed) binds the same site as YYDRxG antibodies but with a different binding mode, although they use similar approach angles that can compete with ACE2.

### Structural studies of group-2 RBD broadly neutralizing antibodies

In our previous study of monoclonal antibodies from COVID-19 recovered-vaccinated donors ^34^, the group-2 antibodies exhibited little or no competition with receptor binding site (RBS) antibodies. Group-2 antibodies showed impressive neutralization breadth against sarbecoviruses including SARS-CoV-2, SARS-CoV-1, Pang17, WIV1, and SHC014 ^34^. Here we show that group 2 antibodies neutralize all SARS-CoV-2 variants to date, including Wuhan, early variants, and Omicron subvariants BA.1, BA.2, BA.5, BQ.1.1, and XBB.1.5 (Table S1). To understand the molecular basis of these broadly neutralizing antibodies, we determined crystal structures of SARS-CoV-2 RBD in complex with three group 2 antibodies, CC25.4, CC25.56 and CC25.43, at resolutions of 1.79 Å, 2.84 Å, and 2.71 Å, respectively (Figure 5 and Table S3). The crystal structures revealed that all three antibodies target a cryptic region on the RBD immediately below the ridge region (Figure 5A). This site has been referred to as ‘site V’ ^31^, ‘left flank’ ^52^, or ‘E3’ ^19^ in previous studies, and does not overlap with the RBS (Figure 5A) or compete with ACE2 binding (Figures S2A and Figure S3).

**Figure 5.**
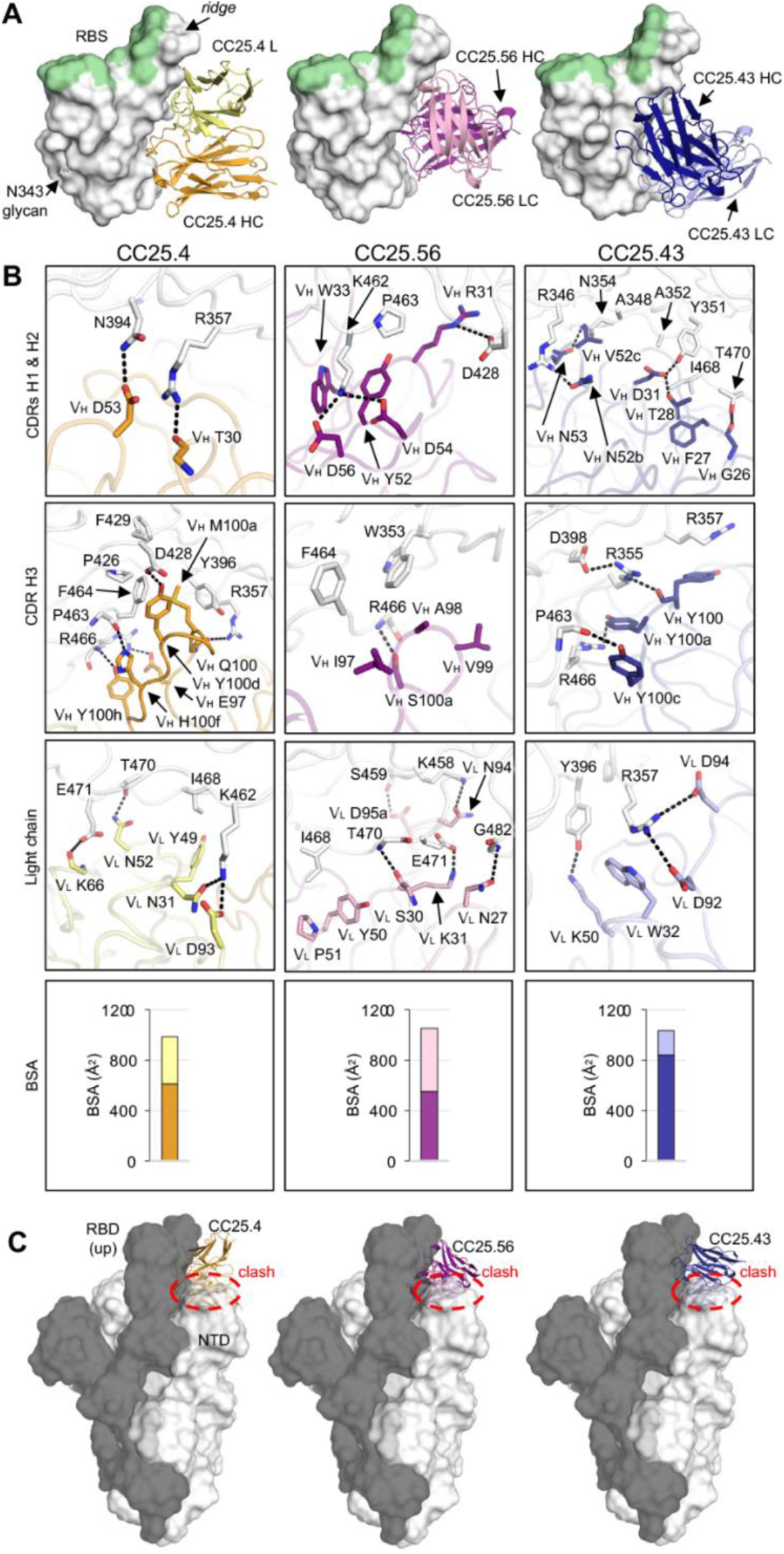
Crystal structures of SARS-CoV-2 RBD in complex with site V antibodies. The SARS-CoV-2 RBD is shown in white, while the receptor binding site (RBS) is highlighted in pale green throughout all the figures. For clarity, only the variable domains of the antibodies are shown in all figures. **A.** Crystal structures of SARS-CoV-2 RBD in complex with site V antibodies. The RBD- N343 N-glycan is shown as sticks. Heavy and light chains of CC25.4 are in orange and yellow, while those of CC25.56 in dark and light pink, and CC25.43 in dark and light blue, respectively. **B.** Detailed interactions between SARS-CoV-2 RBD and the antibodies. Hydrogen bonds and salt bridges are indicated by dashed lines. Surface area of SARS-CoV-2 buried by heavy and light chains of each antibody are shown in the bottom panels. **C.** Models of antibody/RBD structures superimposed onto SARS-CoV-2 spike structures with one-up RBD (PDB 7KJ5). The spike protomer with an up-RBD is shown in grey, while the other two protomers are shown in white. Binding to up-RBDs, site V antibodies would clash with the NTD of the adjacent protomer (indicated with red circles).

All three site V antibodies target the RBD using all six CDRs of their heavy and light chains (Figure 5A, 5B and Figure S4). For CC25.4 CDRs H1 and H2, V_H_ T30 hydrogen bonds with RBD-R357, and V_H_ D53 with RBD-N394. CDR H3 interacts extensively with the RBD (Figure 5B). For example, V_H_ E97 makes a salt bridge with RBD-R466, and V_H_ M100a and Y100d form hydrophobic and aromatic interactions with RBD-Y396, F429, P426, and F464. For the light chain, V_L_ N31 and D93 engage RBD-K462 through a hydrogen bond and salt bridge and other HBs are made with V_L_ N52 and K96 to the RBD (Figure 5B). For CC25.43, the heavy chain dominates the interaction with RBD and is responsible for 82% of BSA. All three HCDR loops interact with RBD. RBD-R357 forms a CH-pi bond with V_H_ Y100, whose main chain interacts with RBD-R355, which in turn forms a salt bridge with D398. RBD-R357 is also clamped by salt bridges to two acidic residues in the light chain, V_L_ D92, and D94. In CC25.56, CDRs H1 and H2 engage in extensive polar interactions with the RBD. V_H_ R31 forms a salt bridge with RBD-D428, and K462 forms two salt bridges with V_H_ D54 and D56. The hydrophobic tip of CDR H3 is comprised of V_H_ I97, A98, and V99, which interact with RBD-W353 and F464. The light chain also forms extensive interactions with the RBD.

RBDs on the SARS-CoV-2 S-protein flip between up and down conformations, where an RBD in the up conformation can engage with the host receptor ACE2, expose more epitope area, and also elicit antibodies specific to up-only conformations. In either up or down-conformation, the S-protein can retain an intact pre-fusion state. However, our structures showed that the site V antibodies target a cryptic site on the RBD. Antibody binding to this epitope would clash with N-terminal domain (NTD) in the adjacent protomer of the S trimer, even if the RBD is in an up conformation when modeled on a pre-fusion SARS-CoV-2 S structure (Figure 5C). This observation suggests that binding of these site V antibodies may result in a conformational rearrangement of the RBD relative to the NTD in the S trimer in the pre-fusion state that could possibly affect viral entry.

Most SARS-CoV-2 RBD antibodies target three regions: the RBS, the CR3022 site, and the S309 site (Figure 6A). Antibodies targeting the RBS generally exhibit higher frequency and higher neutralization potency due to direct competition with receptor ^29^. Almost all commercially available therapeutic neutralizing antibodies (except for Sotrovimab) for COVID-19 treatment target the RBS. However, the RBS is highly variable among sarbecoviruses and SARS-CoV-2 variants (Figures 6A). In fact, all of these therapeutic antibodies have now been evaded by at least one SARS-CoV-2 variant. Here we show that the epitope region of CC25.4, CC25.43, and CC25.56 is highly conserved among sarbecoviruses (Figures 6A-B). Indeed, epitope residues of CC25.4 and CC25.56 are 100% conserved among all SARS-CoV-2 variants to date, while only one mutation (R346T) in BQ.1.1 and XBB.1.5 is located in the CC25.43 epitope (Figure 6B). The high conservation of site V explains the observation that antibodies that target this region largely retain neutralization activity against SARS-CoV-2 variants and other sarbecoviruses (Figure 1B and Table S1) ^34^. In addition, in contrast to some other sites accommodating public antibodies, such as RBS-A targeted by IGHV3-53 and IGHV1-58 antibodies ^6, 53^, RBS-B by IGHV1-2 antibodies ^6^, RBS-D by IGHV2-5 antibodies ^6^, and S2 stem by IGHV1-46 antibodies ^54, 55^, IgBLAST analysis ^56^ showed that antibodies targeting site V are encoded by various germlines and no public antibodies have yet been discovered (Figure 6C).

**Figure 6.**
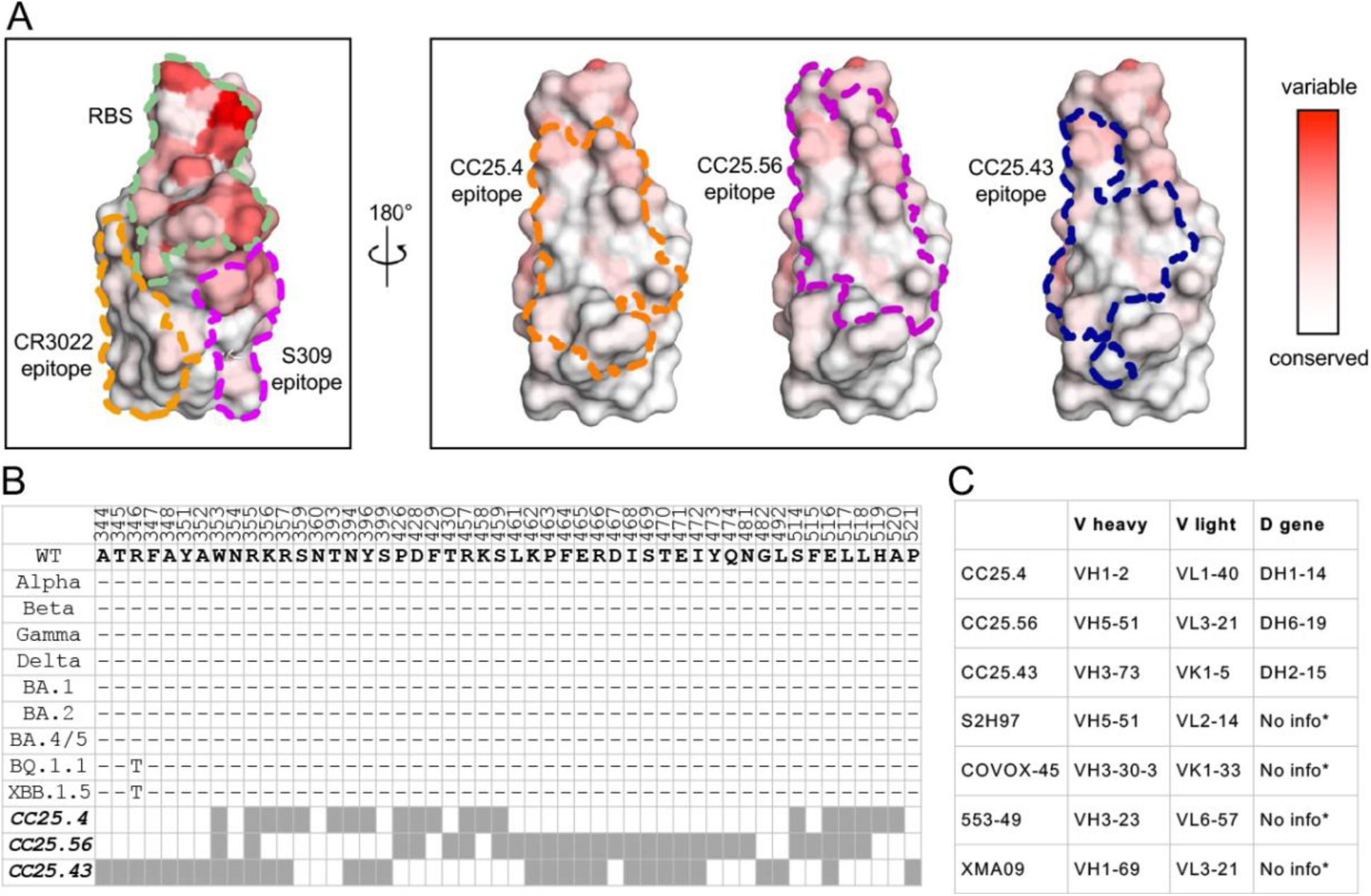
CC25.4, and CC25.56, and CC25.43, target a highly conserved site on the spike RBD. **A.** Locations of the receptor binding site (RBS, pale green) and antibody epitopes (orange, magenta, blue) are indicated by dashed lines [defined as RBD residues with BSA > 0 Å^2^ as calculated by PISA]. A white-red spectrum is used to represent the conservation of each residue of sarbecoviruses including SARS-CoV-2 VOCs, SARS- CoV-1, etc. ^70^ **B.** Sequence alignment of epitope residues of CC25.4, CC25.56, and CC25.43. Identical residues of each variant to the wild-type SARS-CoV-2 are represented by a dash ‘-’. Epitope residues for each antibody are represented as grey boxes. **C.** Putative germline genes encoding site V antibodies as predicted by IgBLAST ^56^. * No info: no D gene or nucleotide sequence information was found for the previously published antibodies S2H97, COVOX-45, 553-49, and XMA09.

To further determine the epitopes recognized by group 2 RBD bnAbs in the soluble spike, we employed single-particle negative stain electron microscopy to image the complexes of SARS-CoV-2 spike and bnAb Fabs (Figure 7). NsEM complexes revealed Fab-induced SARS-CoV-2 spike destabilization with CC25.43 resulting into 100% dimer particles while CC25.4 and CC25.56 showed an approximate 50/50 mix of spike dimer vs. trimer (Figure 7A-C). Particles were unable to converge in 3D due to heterogeneity; therefore, we were unable to produce 3D maps of the spike-antibody complexes. Notably, from the CC25.56 complex, we observed initial binding of antibody to the spike trimers, which quickly dissociated into dimers. Consistent with our crystallography structures, the EM studies revealed that group 2 RBD bnAbs bind to a cryptic face of RBD to destabilize the spike trimer. The findings are suggestive of antibody mediated spike destabilization as putative mechanism of neutralization for group 2 RBD bnAbs.

**Figure 7.**
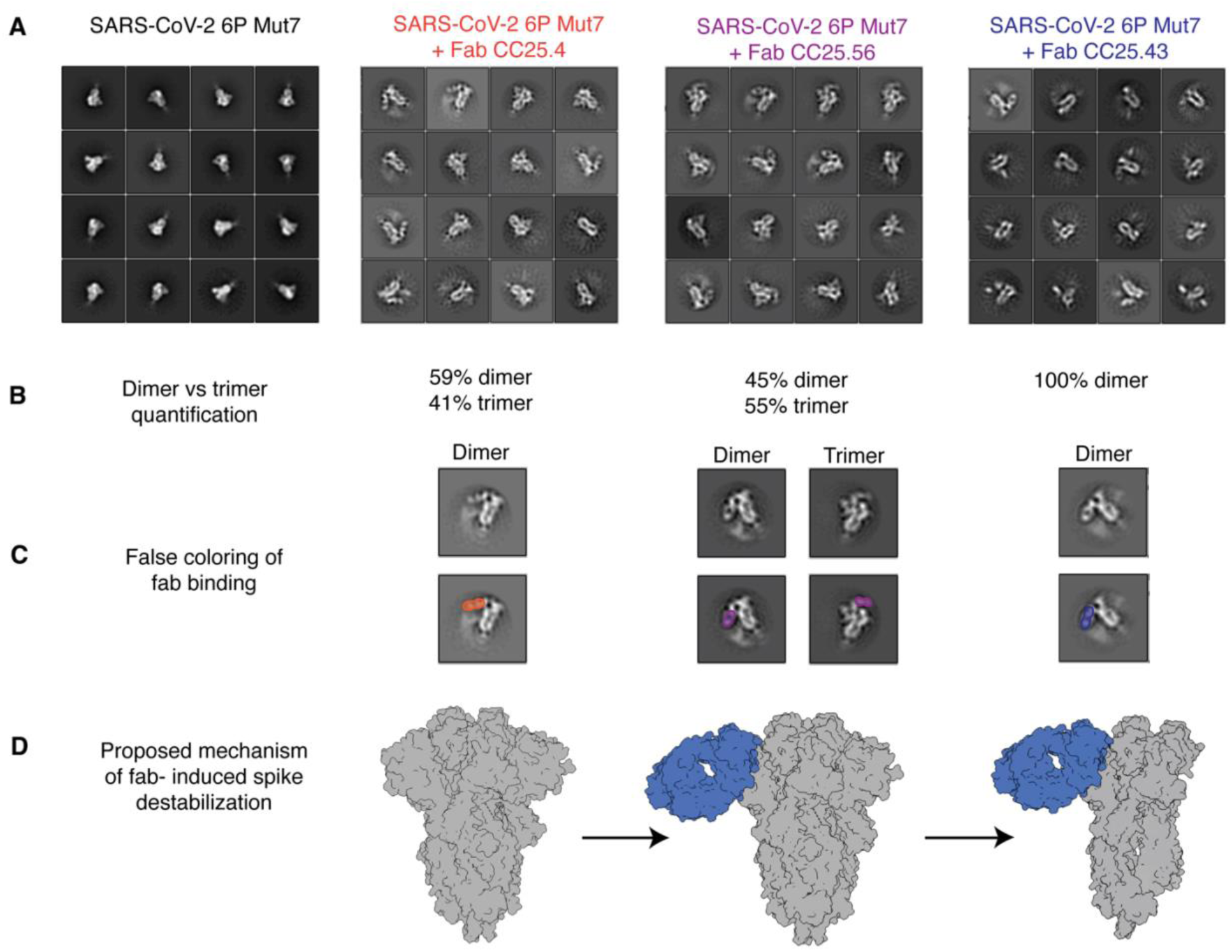
Group 2 RBD bnAbs destabilize the SARS-CoV-2 soluble spike. **A.** Representative 2D classifications of SARS-CoV-2 6P Mut7 spike alone and SARS- CoV-2 6P Mut7 spike in complex with group 2 bnAbs, CC25.4, CC25.56, and CC25.43. **B.** Dimer and trimer proportions were calculated from particle counts within each 2D class. **C.** bnAb binding to the spike is highlighted using false coloring. BnAb binding to trimer was only observed with CC25.56. CC25.4 is colored orange, CC25.56 is colored purple, and CC25.43 is colored blue. **D.** Proposed model of spike degradation induced by bnAb binding. Spike model taken from PDB (6VXX) ^71^. BnAb positioning is approximated based on 2D classes and is colored blue.

## Discussion

The emergence of antibody escape variants of the SARS-CoV-2 Omicron lineage has led to an urgent search for new bnAbs that target conserved spike epitopes. These bnAbs could be used in antibody-mediated prophylaxis or therapy and serve as templates for broad coronavirus vaccine strategies. In this study, we investigated the molecular basis of bnAbs that target conserved spike RBD sites, providing detailed information on the sites that resist virus escape. Our findings will aid in the design of pan-sarbecovirus vaccines.

Here, we revealed that YYDRxG antibodies from group 1 RBD bnAbs (i.e., CC25.54, CC84.24, and CC84.2) target the CR3022 conserved site using essentially the same approach angles as one another, consistent with our previous analysis ^48^. The binding of these antibodies to SARS-CoV-2 RBD sterically clashes with ACE2 binding, although the epitope footprint of these antibodies does not overlap with the ACE2 footprint. The structures demonstrate that diverse light chains permit antibody neutralization. With more interactions between the light chain and SARS-CoV-2 RBD compared to CC25.54 and CC84.24, CC84.2 can neutralize VOCs without somatic hypermutation. We also determined the crystal structure of CC25.36, which binds SARS-CoV-2 with virtually the same approach angle as YYDRxG antibodies and neutralized the virus using the same mechanism as those antibodies. Like CC84.2, the CC25.36 light chain has more contacts compared to CC25.54 and CC84.24, and its inferred germline version showed broad neutralization against VOCs. A newly observed binding motif, YYDML, encoded by IGHD3-9 in CC25.36 CDRH3, binds the same site as the YYDRxG motif but with distinct interactions, which could potentially be a shared motif targeting the CR3022 site. Homology sequence search revealed other potent SARS-CoV-2 RBD Abs, such as COV2-2258 ^8^, C531 ^50^, and C2179 ^51^, share the YYDIL or YYDLL sequences in CDRH3 encoded by IGHD3-9. Further structural studies are needed to determine how similar the binding modes of these antibodies are to that of CC25.36, particularly comparisons of the interaction of YYDIL and YYDLL motifs and the YYDML motif of CC25.36. Overall, multiple antibody germline solutions of group 1 bnAbs converge to recognize a common conserved RBD bnAb site typified by class 4 epitope targeting mAb, CR3022.

In terms of group 1 RBD bnAb B cell precursors, immunogen design strategies could take advantage of the germline-encoded CDRH3 features for vaccine targeting that are described here ^57–61^. One potential challenge is that the immunodominant class 1 and 2 RBS nAbs responses may compete with the group 1 RBD bnAb responses. Accordingly, rational immunogen design may seek to effectively mask the RBS directed strain specific nAb responses, while still favorably exposing the RBD group 1 bnAb site to elicit the desired responses.

We also show that group 2 bnAbs, CC25.4, CC25.43, and CC25.56 target another highly conserved region of the RBD, namely site V, with very few mutations in this site in SARS-CoV-2 variants to date. This epitope region is juxtaposed to the neighboring NTD and likely stabilizes both the up and down conformations of the RBD. Structural and functional roles of this site may therefore be the cause of its extremely low variation. A few previously reported human neutralizing antibodies including S2H97 ^31^, COVOX-45 ^52^, 553-49 ^62^, and XMA09 ^63^ also target this site (Figure S3C) and exhibit remarkably broad neutralization. RBD resurfacing strategies to mask the immunodominant B cell epitopes and immunofocus B cell responses to the site V may be rewarding to induce group 2 bnAb responses (refs).

In addition to site V, some other highly conserved sites on SARS-CoV-2 S have been discovered to be targeted by neutralizing antibodies, including the S2 stem helix ^54, 55, 64–66^ and fusion peptide ^67–69^. Although these sites are well conserved, and therefore can bind antibodies with high breadth, the neutralization potency of these antibodies is usually within the medium-to-low range, potentially due to their indirect neutralizing mechanisms or relative inaccessibility that may require conformational breathing or local rearrangements to permit antibody binding. Further affinity maturation may be required to confer higher neutralizing potency to these broad antibodies.

Overall, our study presents a comprehensive molecular characterization of RBD bnAb epitopes, which can potentially serve as blueprints for the design of broad coronavirus vaccines.

## Acknowledgements

This work was supported by National Institutes of Health-(NIH), National Institute of Allergy and Infectious Diseases-(NIAID) awards, R01AI170928 (R.A.) and CHAVD UM1 AI44462 (D.R.B.) and the Bill and Melinda Gates Foundation INV-004923 (I.A.W., D.R.B.). We thank Henry Tien for technical support with the crystallization robot, and Wenli Yu for protein production. We are grateful to the staff of the Stanford Synchrotron Radiation Lightsource (SSRL) beamlines 12-1 and Advanced Photon Source (APS) beamline 23-ID-B and 23-ID-D for assistance. This research used resources of the SSRL, SLAC National Accelerator Laboratory, which is supported by the U.S. Department of Energy, Office of Science, Office of Basic Energy Sciences under Contract No. DE- AC02–76SF00515. The SSRL Structural Molecular Biology Program is supported by the DOE Office of Biological and Environmental Research, and by the National Institutes of Health, National Institute of General Medical Sciences (including P41GM103393). This research also used resources of the Advanced Photon Source, a U.S. Department of Energy (DOE) Office of Science User Facility, operated for the DOE Office of Science by Argonne National Laboratory under Contract No. DE-AC02-06CH11357. Extraordinary facility operations were supported in part by the DOE Office of Science through the National Virtual Biotechnology Laboratory, a consortium of DOE national laboratories focused on the response to COVID-19, with funding provided by the Coronavirus CARES Act.

## Author contributions

G.S., M.Y., H.L., T.C., D.R.B., I.A.W. and R.A. conceived and designed the study. G.S., T.C., W.H., R.M., K.D., P.Z., S.C., N.M., P.Y., F.A., G.A., A.L.V., X.L., M.M. and L.P. performed BLI, virus preparation, neutralization, and characterization of monoclonal antibodies. Y.S. and B.B performed immunogenetic analysis of antibodies. M.Y., H.L., and Z.F. crystallized the antibody-antigen complexes and determined the crystal structures. M.Y. and H.L., collected X-ray data. M.Y., H.L., X.Z. and I.A.W. analyzed the structural data. R.N.L. and J.L.T. conducted the negative stain electron microscopy studies. G.S., M.Y., H.L., T.C., R.N.L., J.L.T., W.H., R.M., K.D., P.Z., S.C., N.M., P.Y., F.A., G.A., A.L.V., X.L., M.M., Z.F., X.Z., L.P., D.N., Y.S., B.B., A.B.W., D.R.B., I.A.W. and R.A. designed the experiments and/or analyzed the data. G.S., M.Y., H.L., T.C., D.R.B., I.A.W. and R.A. wrote the paper, and all authors reviewed and edited the paper.

## Key Resource Table

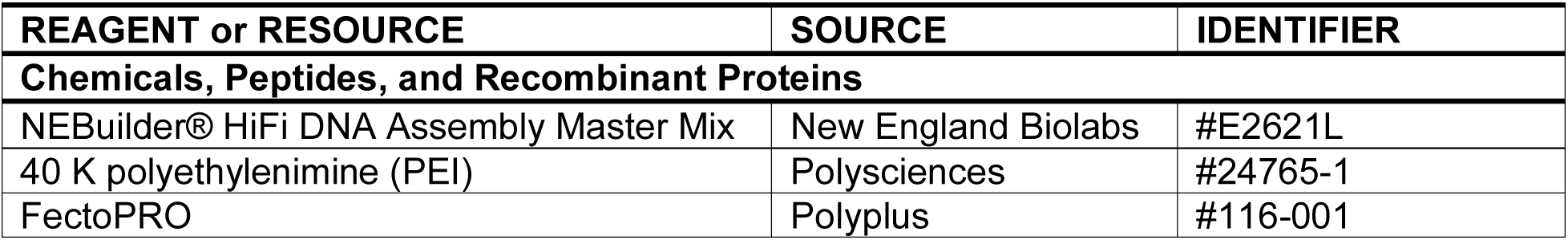
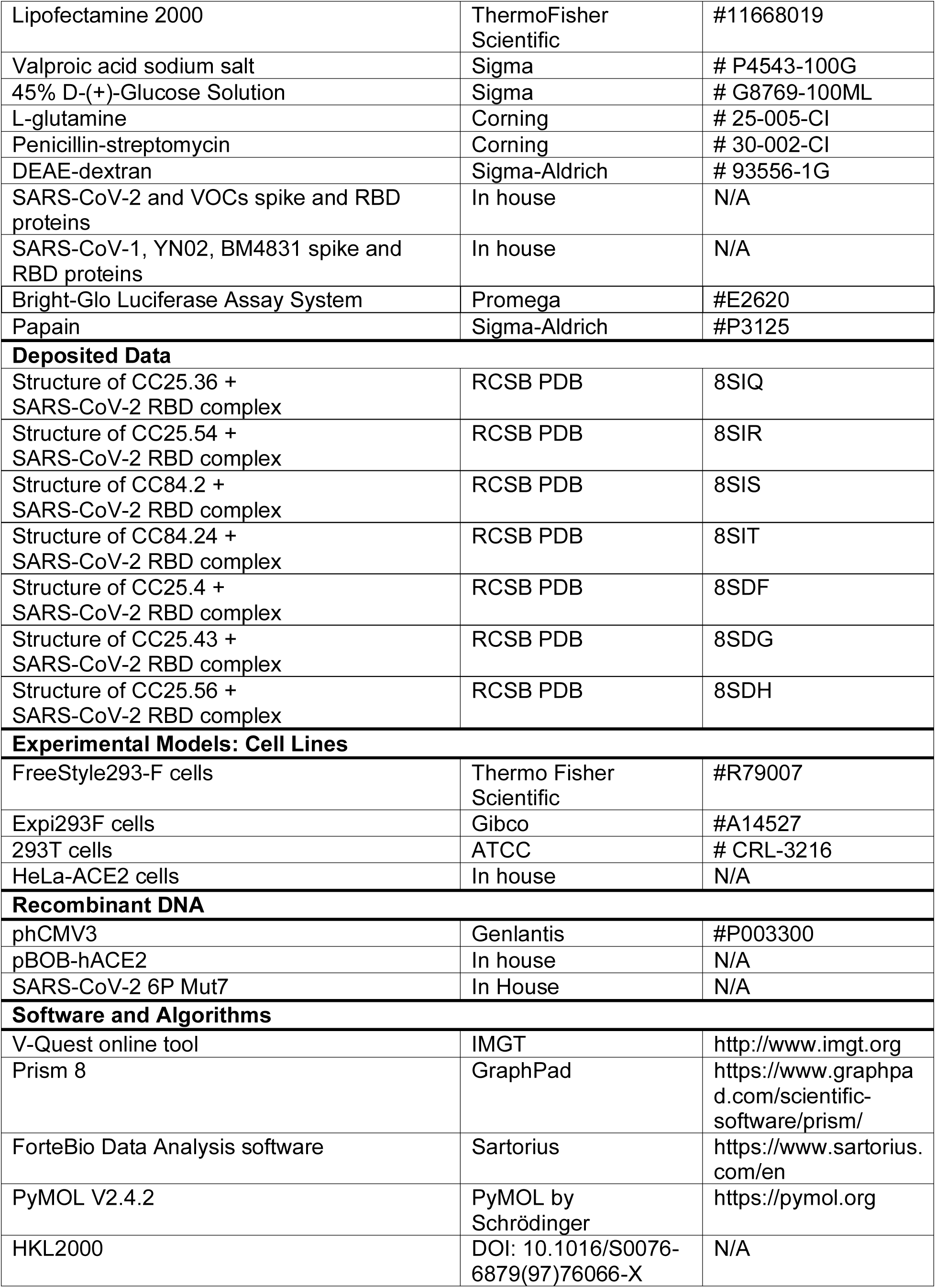
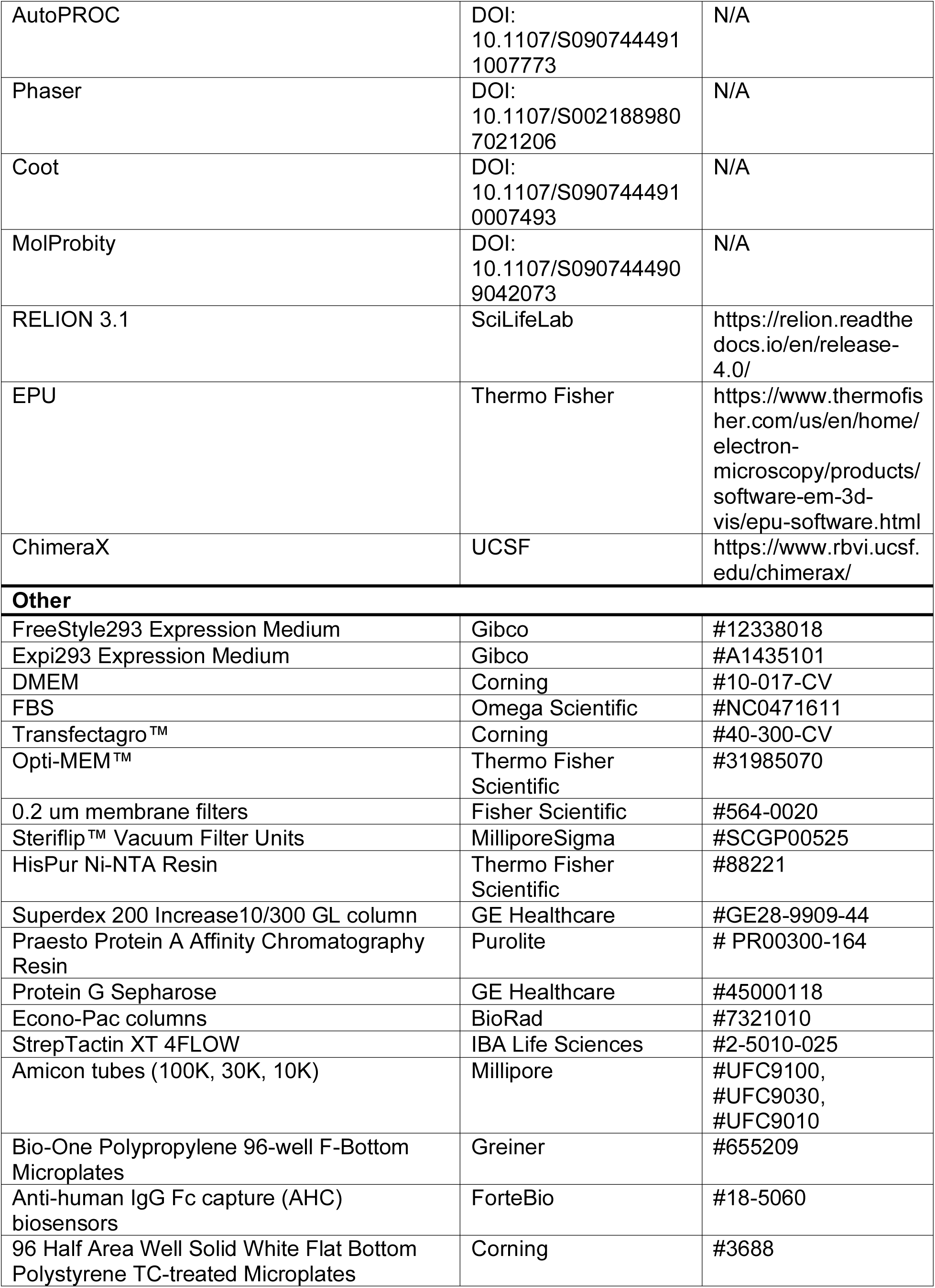
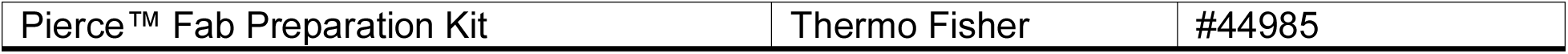

## Resource Availability Lead contact

Further information and requests for resources and reagents should be directed to and will be fulfilled by the lead contact, Raiees Andrabi

## Materials availability

Upon specific request and execution of a material transfer agreement (MTA) from The Scripps Research Institute to the Lead Contact, Antibody plasmids will be made available.

## Data and code availability

The data supporting the findings of this study are available within the published article and summarized in the corresponding tables, figures, and supplemental materials. Inferred germline antibody sequences have been deposited in GenBank under accession numbers XXXX-XXXX. X-ray coordinates and structure factors have been deposited in the RCSB Protein Data Bank under accession codes 8SIQ, 8SIR, 8SIS, 8SIT, 8SDF, 8SDG, and 8SDH.

## MATERIAL AND METHODS

### Expression and purification of spike and RBD proteins

Plasmids coding for the soluble S ectodomain proteins as well as the RBD of human coronaviruses were transfected into Freestyle293F cells (Thermo Fisher R79007). Per 1 L of Freestyle 293F cells, 350 μg of spike/RBD encoding plasmid was mixed with 40 ml transfectagro™ (Corning 40-300-CV) and filtered using 0.22 μm Steriflip™ Sterile Disposable Vacuum Filter Units (MilliporeSigma™ SCGP00525). After filtering, 1.6 mL of 40 K polyethylenimine (Polysciences 24765-1) (1 mg/mL) was added to the plasmid mixture. The resulting solution was gently mixed and then incubate at room temperature for 30 minutes before adding into a 1 L FreeStyle293F cell culture at a concentration of 1 x 10^6^ cells ml^−1^. Four days after transfection, the cell cultures were centrifuged at 2500 x g for 15 minutes and the resulting supernatants were filtered through 0.2 μm membrane filters (Fisher Scientific 564-0020). Cultures were then stored in glass bottles at 4°C before purification. Filtered supernatants containing His-tagged proteins were passed slowly through HisPur Ni-NTA Resin (Thermo Fisher 88221) beads in columns, washed with three bead volumes of wash buffer (25mM Imidazole, pH 7.4) to eliminate non specific binding, and then slowly eluted with 25 ml of elution buffer (250mM Imidazole, pH 7.4). Eluted proteins were buffer exchanged into PBS and then concentrated down using Amicon® 100 kDa or 10 kDa Ultra-15 Centrifugal Filter Units (Merck Millipore UFC9100 & UFC9010), respectively, for spike and RBD proteins. The concentrated proteins were then further purified through size-exclusion chromatography using a Superdex 200 Increase 10/300 GL column (Sigma-Aldrich GE28-9909-44). Selected fractions resulting from the size-exclusion run were pooled together and concentrated again for later use.

### Expression and purification of mAbs

For the expression of mAbs, HC and LC variable regions were cloned into expression vectors with corresponding constant regions. 12 μg plasmid of each chain were mixed into 3 ml Opti-MEM (Gibco 31985070), followed by adding 24 μl FectoPRO transfection reagent (116-040; Polyplus). After incubating at room temperature for 10 min, the mixture was gently added into Expi293F cells (Thermo Fisher Scientific A14527) at a concentration of 2.8 million cells ml^−1^ in final expression volumes of 30 ml. 24 h after transfection, 300 μl sodium valproic acid (300mM) and 275 μl 45% D-(+)-Glucose Solution, (Sigma G8769-100ML) were added. Five days after transfection, the cell cultures were centrifuged at 2500 x g for 15 minutes and the resulting supernatants were filtered using 0.22 μm Steriflip™ Sterile Disposable Vacuum Filter Units (MilliporeSigma™ SCGP00525). The filtered cell culture supernatants were incubated overnight at 4 °C with 0.5 ml 1:1 solution of Praesto Protein A Affinity Chromatography Resin (Purolite PR00300-164) and Protein G Sepharose (Cytiva GE17-0618-01). The solution was then loaded into an Econo-Pac column (Bio-Rad Laboratories 7321010) and washed with 1 column volume of PBS. The mAbs were then eluted with 10 ml 0.2M citric acid (pH 3) and immediately neutralized using 1 ml 2M Tris base (pH 9). The eluted proteins were buffer exchanged into PBS and then concentrated using Amicon® 30 kDa Ultra-15 Centrifugal Filter Units (Merck Millipore UFC9030).

### Production of proteins for the BioLayer Interferometry (BLI) competition analysis

Expression and purification of the SARS-CoV-2 spike protein were done as described previously ^28^. Briefly, the ectodomain (residues 14-1213) with R682G / R683G / R685G / K986P / V987P mutations of the SARS-CoV-2 spike protein (GenBank: QHD43416.1) was cloned into a customized pFastBac vector ^72^. The spike ectodomain constructs were fused with an N-terminal gp67 signal peptide and a C-terminal BirA biotinylation site, thrombin cleavage site, trimerization domain, and His6 tag. Recombinant bacmid DNA was generated using the Bac-to-Bac system (Life Technologies). Baculovirus was generated by transfecting purified bacmid DNA into Sf9 cells using FuGENE HD (Promega), and subsequently used to infect suspension cultures of High Five cells (Life Technologies) at an MOI of 5 to 10. Infected High Five cells were incubated at 28 °C with shaking at 110 rpm for 72 h for protein expression. The supernatant was then concentrated using a Centramate cassette (30 kDa MW cutoff, Pall Corporation). SARS-CoV-2 Spike were purified by Ni-NTA, followed by size exclusion chromatography, and then buffer exchanged into PBS. Expression and purification of the N-terminal peptidase domain of human ACE2 (residues 19 to 615, GenBank: BAB40370.1) was described previously ^73^. ACE2 was cloned into phCMV3 vector and fused with a C-terminal Fc tag. The plasmids were transiently transfected into Expi293F cells using ExpiFectamine™ 293 Reagent (Thermo Fisher Scientific) according to the manufacturer’s instructions. The supernatant was collected at 7 days post-transfection. Fc-tagged ACE2 protein was then purified with a Protein A column (GE Healthcare) followed by size exclusion chromatography.

### BioLayer Interferometry binding assay

Using an Octet RED384 instrument, BLI binding experiments were performed with Anti Human IgG Fc (AHC) biosensors (Sartorius 18-5060). Using Octet buffer (PBS with 0.1% Tween20), mAbs were diluted to 10 μg ml^−1^, while spike and RBD proteins were diluted to 100 nM and 275 nM, respectively. Samples were transferred to black Polypropylene 96-well F-Bottom Microplates (Greiner 655209) for BLI experimentation. The hydrated biosensors first captured the antibodies for 60 s, and then transferred to Octet buffer for 60 s to provide the baseline. The biosensors with captured antibodies were then introduced to the wells containing the viral proteins for 120 s to measure association responses, and into Octet buffer for 240 s to measure disassociation responses. Results from the experiment were analyzed with ForteBio Data Analysis software (version 12) for curve correction and fitting into a 1:1 binding mode. The binding response and KD values were calculated.

For the competition assays of antibodies with ACE2 receptor, Ni-NTA biosensors were used. In brief, the assay had five steps: 1) baseline: 60 s with 1× kinetics buffer; 2) loading: 360 s with 20 μg/mL, His6-tagged SARS-CoV-2 spike protein; 3) baseline: 60 s with 1× kinetics buffer; 4) first association: 360 s with Fabs (2 µM, or buffer only as a control); and 5) second association: 360 s with human ACE2-Fc (200 nM).

### Pseudovirus production

The spike proteins of each tested virus were cloned into a plasmid expression vector with each protein’s endoplasmic reticulum retrieval signal removed. These plasmids were co-transfected with MLV (murine leukemia virus)-CMV (cytomegalovirus) luciferase and MLV Gag/Pol plasmids into HEK-293T cells (ATCC CRL-3216) using Lipofectamine 2000 (Thermo Fisher 11668019) transfection reagent following the manufacturer’s recommended protocol. 16 hours after transfection, cell media was replaced with fresh warm cell media (DMEM (Corning 10-017-CV) with 10% FBS (Omega Scientific NC0471611), 1% L-glutamine (Corning 25-005-CI), and 1% penicillin-streptomycin (Corning 30-002-CI)). 48 hours after transfection, supernatant was collected and filtered using 0.22 μm Steriflip™ Sterile Disposable Vacuum Filter Units (MilliporeSigma™ SCGP00525) and the resulting pseudoviruses were stored at −80 °C for later use.

### Pseudovirus neutralization assay

We first developed hACE2 expressing cells by transducing hACE2 into HeLa cells (ATCC CCL-2) using a lentivirus system. Cells with stable and high hACE2 expression were selected to be used for the pseudovirus neutralization assay. For the neutralization assay, mAbs were diluted to a starting concentration of 60 μg ml^−1^ in cell media (DMEM (Corning 10-017-CV) with 10% FBS (Omega Scientific NC0471611), 1% L-glutamine (Corning 25- 005-CI), and 1% penicillin-streptomycin (Corning 30-002-CI)) and then three-folds serially diluted. 25 μl of each dilution was added to 96 half-area well plates (Corning 3688) and incubated with 25 μl pseudoviruses per well at 37°C for 1 hour. HeLa-hACE2 cells were diluted in cell media to 2 x 10^5^ cells ml^-1^. 50 μl of diluted cells containing 20 μg ml^-1^ DEAE-dextran (Sigma-Aldrich 93556-1G) were added to each well. After 48 hours of incubation at 37°C, supernatant was removed and HeLa-hACE2 cells were lysed with 60 μl per well of a solution containing luciferase lysis buffer (25 mM Gly-Gly, pH 7.8, 15 mM MgSO4, 4 mM EGTA, 1% Triton X-100) and Bright-Glo (Promega Corporation E2620) at a 1:10 ratio. Wells of lysed cells containing Bright Glo were analyzed for luciferase activity using a luminometer. Each mAb was tested in duplicate and repeated independently. Percent neutralization was determined using the equation below:

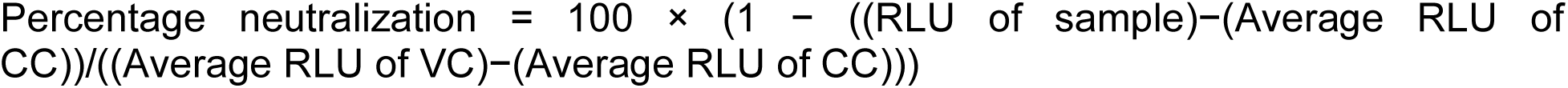

Neutralization percentage was calculated and plotted in Prism 8 (Graph Pad Software) and the IC_50_ antibody titers were determined by fitting a non-linear regression curve and determining the antibody concentration at 50% pseudovirus neutralization.

### Production of proteins for structure analysis

Expression and purification of the SARS-CoV-2 spike receptor-binding domain (RBD) for crystallization were as described previously ^28^. Briefly, the wild-type RBD (residues 333- 529) of the spike (S) proteins was cloned into a customized pFastBac vector ^74^, and fused with an N-terminal gp67 signal peptide and C-terminal His_6_ tag ^28^. The recombinant bacmid DNA was generated using the Bac-to-Bac system (Life Technologies). Baculoviruses were generated by transfecting purified bacmid DNAs into Sf9 cells using FuGENE HD (Promega), and subsequently used to infect suspension cultures of High Five cells (Life Technologies) at an MOI of 5 to 10. Infected High Five cells were incubated at 28 °C with shaking at 110 r.p.m. for 72 h for protein expression. The supernatants were then concentrated using a 10 kDa MW cutoff Centramate cassette (Pall Corporation). The RBD protein was purified by Ni-NTA, followed by size exclusion chromatography, and buffer exchanged into 20 mM Tris-HCl pH 7.4 and 150 mM NaCl.

Fabs used for crystallization were expressed and purified as follows: the plasmids of heavy and light chains were transiently co-transfected into Expi293F cells using FectoPRO transfection reagent (116-040; Polyplus) according to the manufacturer’s instructions. The supernatant was collected at 5 days post-transfection. The Fab was purified with a CaptureSelect CH1-XL Pre-packed Column (Thermo Fisher Scientific) followed by size exclusion chromatography. VSRRLP and VFNQIKP variants of the elbow region was used to reduce the conformational flexibility between the heavy chain constant and variable domains of Fabs CC25.4 and CC25.56, respectively, for crystallization ^75^. For CC25.43, both the heavy and light chains were mutated to facilitate crystal packing. A VFNQIKG mutation was applied to the elbow region of the heavy chain ^75^, and the FG loop of the kappa chain (HQGLSSP) was shortened to QGTTS to facilitate edge-to-edge beta-sheet packing ^76^. Other structures in this paper were obtained from crystallization with unmutated Fabs and RBD.

Fabs used for ns-EM were purified as follows: expressed IgGs were concentrated and digested using the Pierce™ Fab Preparation Kit (Thermo Fisher 44985) following manufacturer instructions. The resulting Fabs were buffer exchanged into PBS and then concentrated down using Amicon® 10 kDa Ultra-15 Centrifugal Filter Units (Merck Millipore UFC9010). Selected fractions resulting from the size-exclusion chromatography were pooled together and concentrated again for later use.

### Crystallization and structural determination

CC25.4/RBD, CC25.36/RBD/CV38-142, CC25.54/RBD, CC25.43/RBD, CC25.56/RBD, CC84.2/RBD, CC84.24/RBD complexes were formed by mixing each of the protein components in an equimolar ratio and incubating overnight at 4°C. The protein complexes were adjusted to 8.6–12 mg/ml and screened for crystallization using the 384 conditions of the JCSG Core Suite (Qiagen) on our robotic CrystalMation system (Rigaku) at Scripps Research. Crystallization trials were set-up by the vapor diffusion method in sitting drops containing 0.1 μl of protein and 0.1 μl of reservoir solution. For the CC25.4/RBD complex, optimized crystals were grown in drops containing 65% MPD and 0.1 M Bicine pH 9.0 at 20°C. Crystals appeared on day 28 and were harvested on day 30. Diffraction data were collected at cryogenic temperature (100 K) at beamline 23-ID-B of the Advanced Photon Source (APS) at Argonne National Labs. For the CC25.36/RBD/CV38-142 complex, optimized crystals were grown in drops containing 20% (w/v) PEG-3350 and 0.2 M diammonium citrate at 20°C. Crystals appeared on day 7 and were harvested on day 15. Diffraction data were collected at cryogenic temperature (100 K) at beamline 12-1 of the Stanford Synchrotron Radiation Lightsource (SSRL). For the CC25.54/RBD complex, optimized crystals were grown in drops containing 20% (w/v) PEG-3350 and 0.2 M potassium sodium tartrate pH 7.2 at 20°C. Crystals appeared on day 28 and were harvested on day 45. Diffraction data were collected at cryogenic temperature (100 K) at beamline 23-ID-B of the Advanced Photon Source (APS) at Argonne National Labs. For the CC25.43/RBD complex, optimized crystals were grown in drops containing 1.0 M Li- chloride, 10% PEG-6000, and 0.1 M citric acid pH 4.0 at 20°C. Crystals appeared on day 7 and were harvested on day 15 by soaking in reservoir solution supplemented with 20% (v/v) ethylene glycol. Diffraction data were collected at cryogenic temperature (100 K) at beamline 23-ID-B of the APS. The B-values of the RBD molecules in the CC25.43/RBD complex structure are higher than the overall average B-values because a large region of the RBDs is exposed in the crystal with lack of stabilization from crystal packing. For the CC25.56/RBD complex, optimized crystals were grown in drops containing 10% ethylene glycol (v/v), 0.11 M MgCl_2_, and 16% polyethylene glycol 3350 (w/v) at 20°C. Crystals appeared on day 7 and were harvested on day 15 with no additional cryoprotectant. Diffraction data were collected at cryogenic temperature (100 K) at beamline 23-ID-D of the APS. For the CC84.2/RBD complex, optimized crystals were grown in drops containing 20% (w/v) PEG-3000 and 0.1 M sodium citrate pH 5.5 at 20°C. Crystals appeared on day 21 and were harvested on day 45. Diffraction data were collected at cryogenic temperature (100 K) at beamline 23-ID-B of the APS. For the CC84.24/RBD complex, optimized crystals were grown in drops containing 20% (w/v) PEG-3000 and 0.1 M Sodium citrate pH 5.5 at 20°C. Crystals appeared on day 21 and were harvested on day 45. Diffraction data were collected at cryogenic temperature (100 K) at beamline 23-ID-B of the APS. Diffraction data were processed with either HKL2000 (PubMed: 27754618) or AutoPROC ^77^. Structures were solved by molecular replacement using PHASER ^78^. Iterative model building and refinement were carried out in COOT ^79^ and PHENIX ^80^, respectively. Epitope and paratope residues, as well as their interactions, were identified by accessing PISA at the European Bioinformatics Institute (http://www.ebi.ac.uk/pdbe/prot_int/pistart.html) ^81^.

### Negative stain electron microscopy

SARS-CoV-2 6P Mut7 and monoclonal bnAbs were complexed at a 3:1 molar ratio (Fab:spike) for one hour at room temperature and SEC purified using a Superose 6 Increase 10/300 GL column in an AKTA Pure system. nsEM grids were made by depositing 3 µl of complex at a dilution of ∼0.03 mg/ml and stained with 2% uranyl formate for 90 seconds. Grids were imaged using a Thermo Fisher Falcon 4i Direct Electron Detector 4K x 4K camera on a Thermo Fisher Glacios (200 kEV, 73kx mag) and micrographs were processed in Relion 3.1 ^82^. Particles from raw micrographs were picked using a Laplacian-of-Gaussian spatial filter and classified into 2D classes. Cartoon figures were made in ChimeraX ^83^.

### Statistical analysis

Statistical analysis was performed using Graph Pad Prism 8, Graph Pad Software, San Diego, California, USA. IC_50_ neutralization titers or BLI binding responses were compared using the non-parametric unpaired Mann-Whitney-U test. Data were considered statistically significant when p < 0.05.

**Figure S1.**
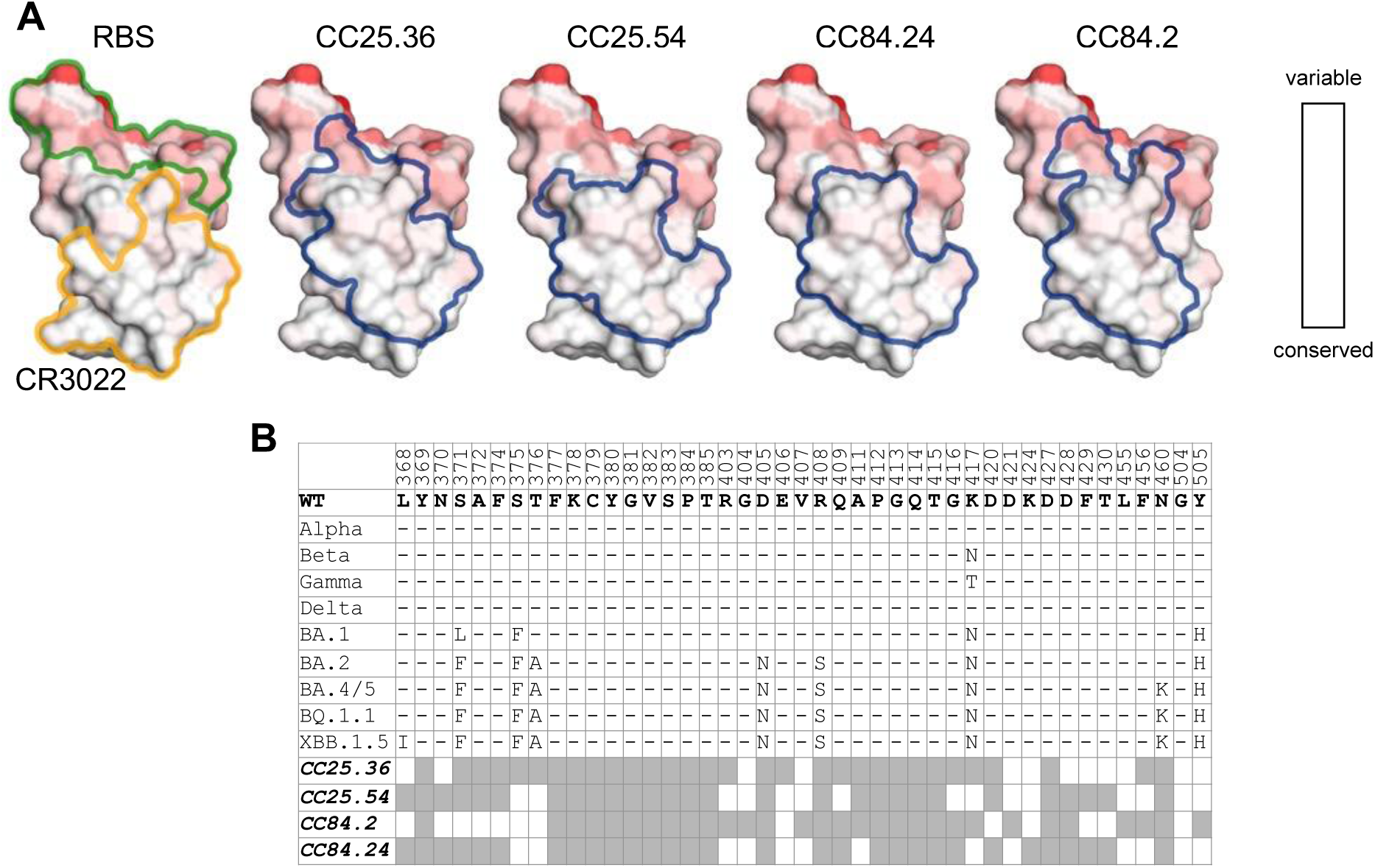
Group 1 RBD bnAb target a relatively conserved site targeted by class 4 bnAbs. **A.** Locations of the receptor binding site (RBS, pale green) and RBD class 4 site CR3022 antibody epitope (orange) are indicated by outlines [defined as RBD residues with buried surface area (BSA) > 0 Å^2^ as calculated by PISA]. A white-red spectrum is used to represent the conservation of each residue of sarbecoviruses including SARS-CoV-2 VOCs, SARS-CoV-1, etc. as in the Figure 6A. **B.** Sequence alignment of epitope residues of group 1 RBD bnAbs, CC25.36, CC25.54, CC84.2 and CC84.24. Identical residues of each variant to the wild-type SARS-CoV-2 are represented by a dash ‘-’. Epitope residues for each antibody are represented as grey boxes.

**Figure S2.**
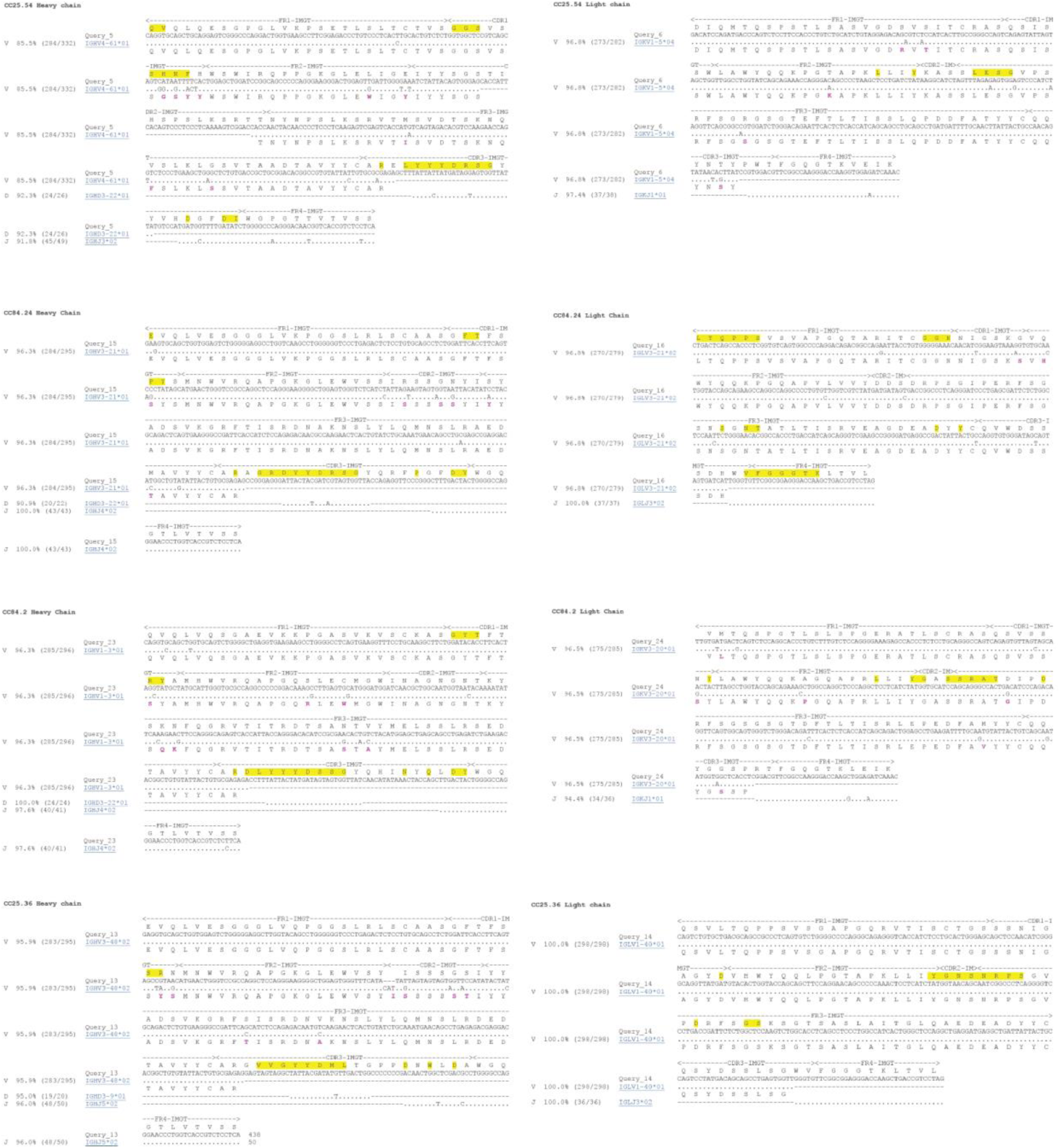
Alignment of group 1 RBD broadly neutralizing antibodies and their putative germline sequences. Paratope residues [defined as buried surface area (BSA) > 0 Å^2^ as calculated by PISA ^81^ of group 1 RBD bnAbs, CC25.54, CC84.24, CC84.2 and CC25.36 are highlighted with yellow boxes. Germline residues that have been somatically mutated as calculated by IgBLAST ^56^ are highlighted in purple.

**Figure S3.**
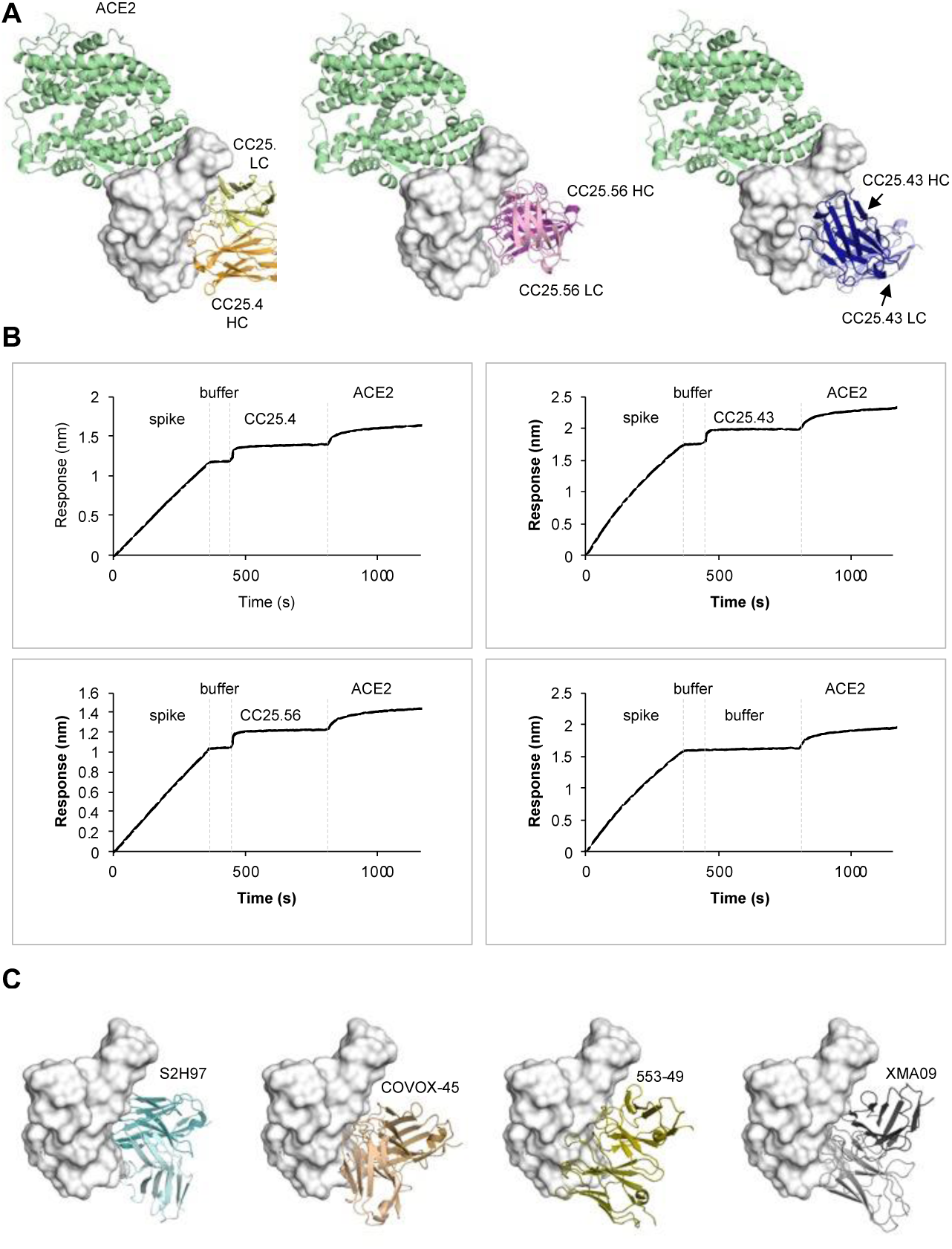
Structures of antibodies targeting site V of SARS-CoV-2 RBD. The SARS-CoV-2 RBD is shown in white. **A.** The structure of SARS-CoV-2 RBD in complex with human receptor ACE2 (PDB 6M0J) was superimposed onto the structures of RBD in complex with site-V antibodies determined in this study. ACE2 is in pale green. Heavy and light chains of CC25.4 are in orange and yellow, while those of CC25.56 in dark and light pink, and CC25.43 in dark and light blue, respectively. **B.** Biolayer interferometry assay of ACE2 binding to SARS-CoV-2 S protein in the presence of site V targeting Fabs. **C.** Comparison with previously reported site-V antibodies. Antibodies S2H97 (PDB 7M7W), COVOX-45 (PDB 7ORA), 553-49 (PDB 7WOG), and XMA09 (PDB 7WHZ) are shown in teal, sand, olive, and grey, respectively (heavy and light chains in dark and light colors).

**Figure S4.**
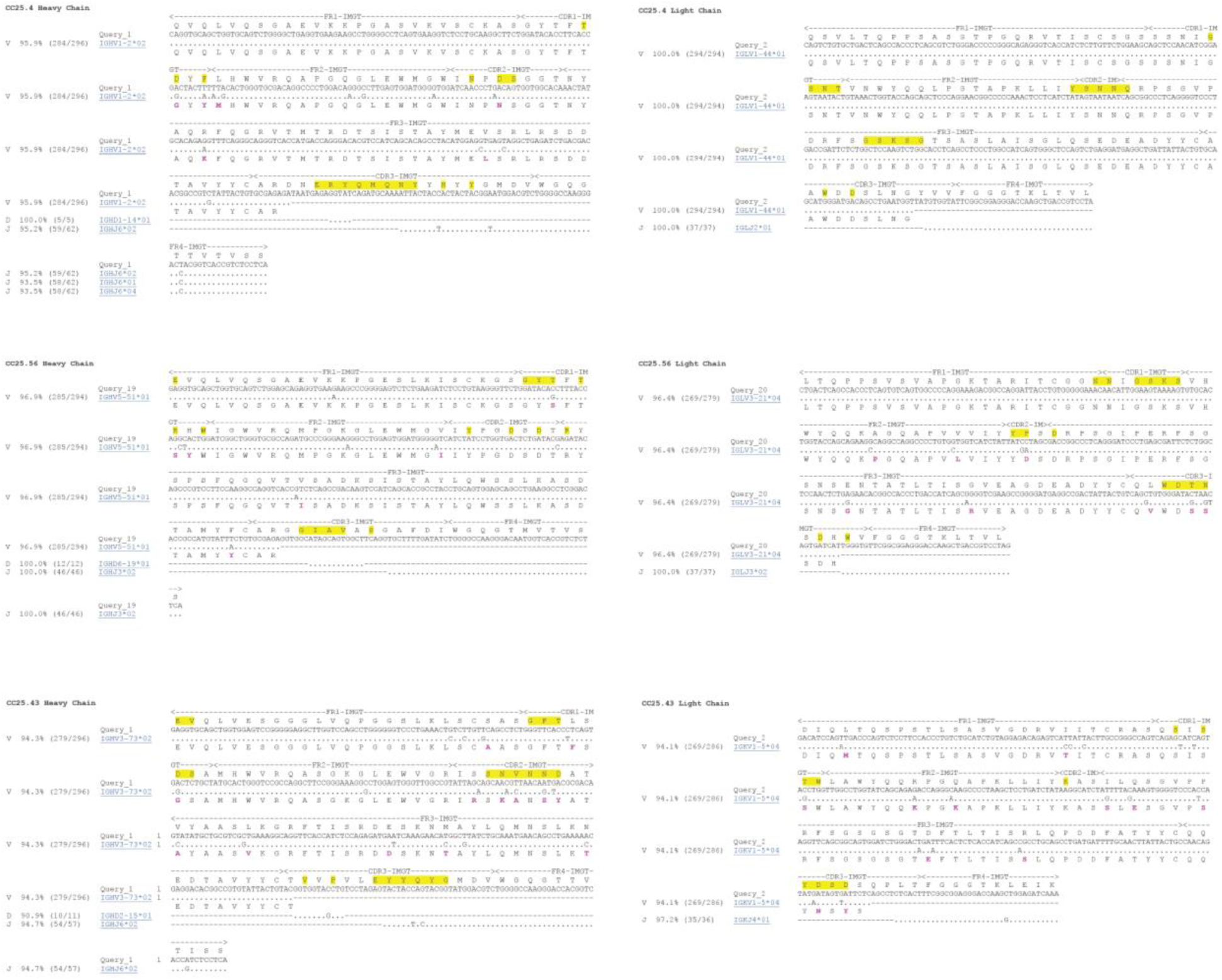
Alignment of group 1 and group 2 RBD broadly neutralizing antibodies and their putative germline sequences. Paratope residues [defined as buried surface area (BSA) > 0 Å^2^ as calculated by PISA ^81^ of group 2 RBD bnAbs, CC25.4, CC25.56 and CC25.43 are highlighted with yellow boxes. Germline residues that have been somatically mutated as calculated by IgBLAST ^56^ are highlighted in purple.

**Table S1.**
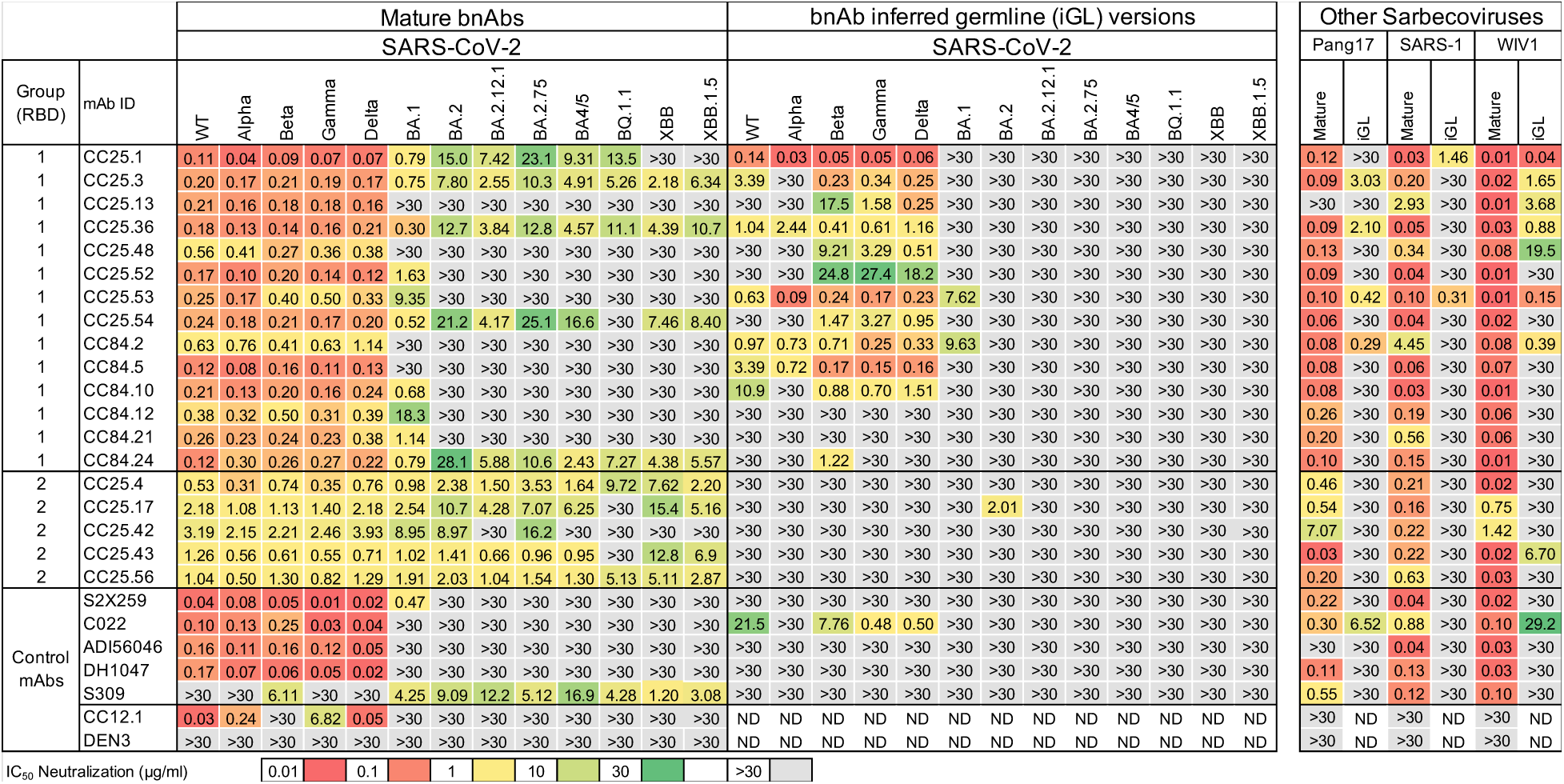
IC_50_ Neutralization of group 1 and 2 bnAbs and their inferred germline antibodies. Experimentally determined neutralization IC_50_s of the mature and inferred germline (iGL) versions of 14 Group 1 and 5 Group 2 RBD bnAbs tested with pseudotyped versions of SARS-CoV-2 (Wuhan), 12 SARS-CoV-2 Variants of Concern (B.1.1.7 (Alpha), B.1.351 (Beta), P.1 (Gamma), Delta (B.1.617.2), BA.1 (Omicron), BA.2, BA.2.12.1, BA.2.75, BA.4/5, BQ.1.1, XBB, XBB.1.5), and 3 other sarbecoviruses (Pang17 (Clade 1b), SARS-CoV-1 (Clade 1a), and WIV1 (Clade 1a)). Control Abs used were SARS-CoV-2 antibodies S2X259, C022, ADI56046, DH1047, S309, and CC12.1, and the Dengue antibody, DEN3. All experiments were conducted independently twice and verified to produce similar results. ND = Not Determined.

**Table S2.**
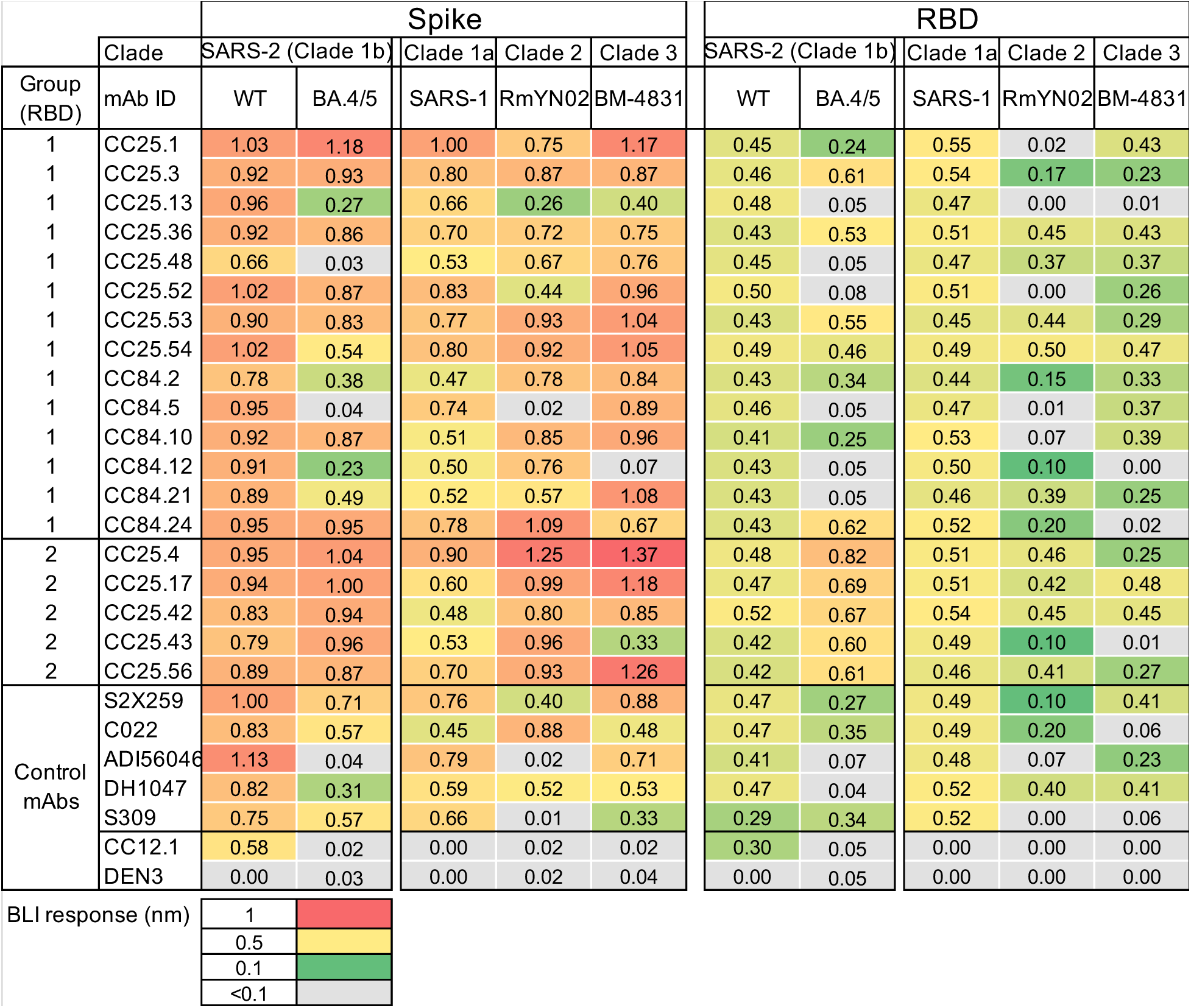
BLI binding responses. BLI response values (nm) of 14 Group 1 and 5 Group 2 RBD bnAbs binding to the trimeric spike proteins and monomeric RBD proteins of Clade 1b (SARS-CoV-2 and BA.4/5), Clade 1a (SARS-CoV-1), Clade 2 (RmYN02), and Clade 3 (BM-4831) betacoronaviruses. Control antibodies used were SARS-CoV-2 Abs S2X259, C022, ADI56046, DH1047, S309, and CC12.1, and the Dengue antibody DEN3.

**Table S3.** **X-ray data collection and refinement statistics**

| **Data collection** | CC25.36 Fab + SARS-CoV-2 RBD + CV38-142 Fab | CC25.54 Fab + SARS-CoV-2 RBD | CC84.2 Fab + SARS-CoV-2 RBD | CC84.24 Fab + SARS-CoV-2 RBD |
| --- | --- | --- | --- | --- |
| Beamline | SSRL 12-1 | APS 23-IDB | APS 23-IDB | APS 23-IDB |
| Wavelength (Å) | 0.97946 | 0.97930 | 1.03317 | 1.03317 |
| Space group | *P* 2<sub>1</sub> 2<sub>1</sub> 2<sub>1</sub> | *P* 3<sub>2</sub> 2 1 | *P* 1 2<sub>1</sub> 1 | *P* 2<sub>1</sub> 2<sub>1</sub> 2<sub>1</sub> |
| Unit cell parameters |  |  |  |  |
| a, b, c (Å) | 60.4, 77.0, 266.4 | 135.9, 135.9, 87.1 | 68.7, 80.2, 73.5 | 101.4, 106.0, 247.0 |
| α, β, γ (°) | 90, 90, 90 | 90, 90, 90 | 90, 94.9, 90 | 90, 90, 90 |
| Resolution (Å)<sup>a</sup> | 50.0–2.50 (2.54–2.50) | 50.0–3.30 (3.36–3.30) | 50.0–3.08 (3.15–3.08) | 50.0–2.91 (2.95–2.91) |
| Unique reflections<sup>a</sup> | 38,811 (1,970) | 14,256 (705) | 14,522 (706) | 54,891 (2,362) |
| Redundancy<sup>a</sup> | 3.5 (3.3) | 11.5 (7.1) | 4.2 (4.2) | 6.2 (3.5) |
| Completeness (%)<sup>a</sup> | 87.8 (92.0) | 99.8 (98.7) | 97.9 (97.0) | 92.5 (81.3) |
| <I/σ<sub>I</sub>><sup>a</sup> | 7.9 (1.1) | 8.0 (2.0) | 6.3 (1.0) | 6.9 (1.2) |
| *R*<sub>sym</sub><sup>b</sup> (%)<sup>a</sup> | 15.7 (>100) | 29.8 (97.8) | 20.5 (>100) | 21.4 (76.6) |
| *R*<sub>pim</sub><sup>b</sup> (%)<sup>a</sup> | 8.8 (66.0) | 9.1 (38.1) | 11.2 (63.6) | 8.9 (43.0) |
| CC<sub>1/2</sub><sup>c</sup> (%)<sup>a</sup> | 98.0 (51.5) | 98.8 (32.1) | 99.2 (71.7) | 98.3 (42.1) |
| **Refinement statistics** |  |  |  |  |
| Resolution (Å) | 38.5–2.50 | 48.8–3.30 | 48.0–3.08 | 48.7–2.91 |
| Reflections (work) | 32,649 | 10,644 | 10,876 | 44,227 |
| Reflections (test) | 1,635 | 542 | 563 | 2,275 |
| *R*<sub>cryst</sub><sup>d</sup> / *R*<sub>free</sub><sup>e</sup> (%) | 24.7/28.7 | 20.8/23.6 | 25.6/30.3 | 26.1/30.9 |
| No. of atoms | 7,963 | 4,803 | 4,647 | 18,011 |
| Macromolecules | 7,914 | 4,789 | 4,633 | 17,983 |
| Glycans | 49 | 14 | 14 | 28 |
| Average *B*-value (Å<sup>2</sup>) | 50 | 68 | 78 | 66 |
| Macromolecules | 50 | 68 | 78 | 66 |
| Fab | 51 | 67 | 70 | 70 |
| RBD | 46 | 70 | 96 | 58 |
| Glycans | 48 | 87 | 127 | 67 |
| Wilson *B*-value (Å<sup>2</sup>) | 40 | 53 | 71 | 48 |
| **RMSD from ideal geometry** |  |  |  |  |
| Bond length (Å) | 0.005 | 0.002 | 0.004 | 0.006 |
| Bond angle (°) | 0.76 | 0.57 | 0.69 | 1.04 |
| **Ramachandran statistics (%)** |  |  |  |  |
| Favored | 98.7 | 94.9 | 97.6 | 97.6 |
| Outliers | 0.0 | 0.0 | 0.0 | 0.0 |
| **PDB code** | 8SIQ | 8SIR | 8SIS | 8SIT |

| Data collection | CC25.4 +<br>SARS-CoV-2 RBD | CC25.43 +<br>SARS-CoV-2 RBD | CC25.56 +<br>SARS-CoV-2 RBD |
| --- | --- | --- | --- |
| Beamline | APS23ID-B | APS23ID-B | APS23ID-D |
| Wavelength (Å) | 1.0332 | 1.0332 | 1.0332 |
| Space group | P 1 2<sub>1</sub> 1 | C 1 2 1 | C 1 2 1 |
| Unit cell parameters |  |  |  |
| a, b, c (Å) | 58.2, 143.0, 82.9 | 311.9, 83.8, 74.1 | 171.1, 105.6, 99.0 |
| α, β, γ (°) | 90, 94.1, 90 | 90, 100.8, 90 | 90, 117.3, 90 |
| Resolution (Å)<sup>a</sup> | 82.7-1.79 (1.82-1.79) | 153.2-2.71 (2.76-2.71) | 77.8-2.84 (2.89-2.84) |
| Unique reflections<sup>a</sup> | 126,496 (6,298) | 47,774 (2,451) | 36,671 (1,820) |
| Redundancy<sup>a</sup> | 7.0 (6.5) | 5.2 (5.2) | 4.3 (4.2) |
| Completeness (%)<sup>a</sup> | 99.2 (98.6) | 93.4 (96.8) | 99.1 (98.8) |
| <I/σ<sub>I</sub>><sup>a</sup> | 9.8 (0.7) | 7.8 (1.0) | 6.2 (1.1) |
| R<sub>sym</sub><sup>b</sup> (%)<sup>a</sup> | 9.4 (>100) | 16.5 (>100) | 24.7 (>100) |
| R<sub>pim</sub><sup>b</sup> (%)<sup>a</sup> | 3.8 (>100) | 7.8 (98.4) | 13.6 (96.7) |
| CC<sub>1/2</sub><sup>c</sup> (%)<sup>a</sup> | 99.8 (35.6) | 99.5 (49.5) | 97.1 (36.4) |
| **Refinement statistics** |  |  |  |
| Resolution (Å) | 54.1-1.79 | 72.8-2.71 | 47.7-2.84 |
| Reflections (work) | 125,886 | 47,703 | 36,574 |
| Reflections (test) | 12,469 | 4,945 | 36,05 |
| R<sub>cryst</sub><sup>d</sup> / R<sub>free</sub><sup>e</sup> (%) | 18.6/21.8 | 26.0/30.4 | 24.6/28.3 |
| No. Fab/RBD copies in ASU | 2 | 2 | 2 |
| No. of atoms | 10,521 | 9,536 | 9,606 |
| RBD | 3,149 | 3,006 | 3,118 |
| Fab | 6,528 | 6,530 | 6,447 |
| Ligands<sup>f</sup> | 64 | 0 | 0 |
| Solvent | 780 | 0 | 41 |
| Average B-values (Å<sup>2</sup>) | 43 | 89 | 55 |
| RBD | 42 | 128 | 71 |
| Fab | 44 | 71 | 73 |
| Ligands<sup>f</sup> | 60 | N/A | N/A |
| Solvent | 46 | N/A | 51 |
| Wilson B-value (Å<sup>2</sup>) | 32 | 61 | 55 |
| **RMSD from ideal geometry** |  |  |  |
| Bond length (Å) | 0.010 | 0.004 | 0.004 |
| Bond angle (°) | 1.1 | 0.77 | 0.87 |
| **Ramachandran statistics (%)<sup>g</sup>** |  |  |  |
| Favored | 96.3 | 96.7 | 97.3 |
| Outliers | 0.32 | 0.16 | 0.24 |
| **PDB code** | **8SDF** | **8SDG** | **8SDH** |

<sup>a</sup> Numbers in parentheses refer to the highest resolution shell.
<sup>b</sup> $R_{\text{sym}} = \sum_{hkl} \sum_i |I_{hkl,i} - \langle I_{hkl} \rangle| / \sum_{hkl} \sum_i I_{hkl,i}$ and $R_{\text{pim}} = \sum_{hkl} (1/(n-1))^{1/2} \sum_i |I_{hkl,i} - \langle I_{hkl} \rangle| / \sum_{hkl} \sum_i I_{hkl,i}$ , where $I_{hkl,i}$ is the scaled intensity of the $i^{\text{th}}$ measurement of reflection $h, k, l$ , $\langle I_{hkl} \rangle$ is the average intensity for that reflection, and $n$ is the redundancy.
<sup>c</sup> CC<sub>1/2</sub> = Pearson correlation coefficient between two random half datasets.
<sup>d</sup> $R_{\text{cryst}} = \sum_{hkl} |F_o - F_c| / \sum_{hkl} |F_o| \times 100$ , where $F_o$ and $F_c$ are the observed and calculated structure factors, respectively.
<sup>e</sup> $R_{\text{free}}$ was calculated as for $R_{\text{cryst}}$ , but on a test set comprising 2.5% or 5% of the data excluded from refinement.
<sup>f</sup> Bound ligands are MPD.
<sup>g</sup> From MolProbity<sup>84</sup>.

